# Yeast diversity in open agave fermentations across Mexico

**DOI:** 10.1101/2023.07.02.547337

**Authors:** Porfirio Gallegos-Casillas, Luis F. García-Ortega, Adriana Espinosa-Cantú, Carolina G. Torres-Lagunes, J. Abraham Avelar-Rivas, Adrián Cano-Ricardez, Ángela M. García-Acero, Susana Ruiz-Castro, Mayra Flores-Barraza, Alejandra Castillo, Fernando González-Zozaya, América Delgado-Lemus, Francisco Molina-Freaner, Cuauhtémoc Jacques-Hernández, Antonio Hernández-López, Luis Delaye, Xitlali Aguirre-Dugua, Manuel R. Kirchmayr, Lucía Morales, Eugenio Mancera, Alexander DeLuna

## Abstract

Yeasts are a diverse group of fungal microorganisms that are widely used to produce fermented foods and beverages. In Mexico, open fermentations are used to obtain spirits from agave plants. Despite the prevalence of this traditional practice throughout the country, yeasts have only been isolated and studied from a limited number of distilleries. To systematically describe the diversity of yeast species from open agave fermentations, here we generate the YMX.1.0 culture collection by isolating 4,524 strains from 68 sites in diverse climatic, geographical, and biological contexts. We used MALDI-TOF mass spectrometry for taxonomic classification and a subset of the strains was verified by ITS and D1/D2 sequencing. The most abundant species isolated from all producing regions were *Saccharomyces cerevisiae*, *Pichia kudriavzevii*, *Pichia manshurica*, and *Kluyveromyces marxianus*. Despite the great diversity of environmental conditions and production practices the composition of yeast communities remained largely homogeneous throughout locations and fermentation stages, even if less abundant but commonly occurring yeasts were considered. Furthermore, ITS and D1/D2 sequencing revealed two candidate new species of Saccharomycetales. To explore the intraspecific variation of the yeasts from agave fermentations, we conducted genome sequencing on four isolates of the non-conventional yeast *Kazachstania humilis*. The genomes of these four strains were considerably distinct from other genomes of the same species, suggesting that they belong to a different population. Our work contributes to the understanding and conservation of an open fermentation system of great cultural and economic importance, providing a valuable resource to study the biology and genetic diversity of microorganisms living at the interface of natural and human-associated environments.

**TAKE AWAY:**

- We isolated and identified 4,524 yeast strains from open agave fermentations in Mexico.
- Yeast communities remained largely homogeneous throughout diverse locations.
- *Kazachstania humilis* genomes differed significantly from isolates in other regions of the world.
- We report two candidate new species related to the *Pichia* clade.

## INTRODUCTION

Yeasts are single-celled fungi that have played a significant role in human history. These microorganisms have a wide geographical distribution, in part as a result of their use by humans in the production of fermented foods and beverages; they occur naturally in a variety of climates, ecosystems, and raw fermentation materials such as grapes and cereals (Gallone et al., 2016; Giannakou et al., 2020; Marsit et al., 2017; Molinet & Cubillos, 2020). The first archaeological records of fermented-beverage production are dated to 8,000-3,000 BC in China, Sumeria, Iran, and Egypt (De Chiara et al., 2022; Marsit et al., 2017; Turker, M. 2015). Way before humans knew about their biological nature, yeasts had been handled in fermentations to produce wine, ciders, distilled spirits, bread, cheese, and other products. In addition, Saccharomycetales yeasts are used nowadays in a variety of biotechnological applications, such as bioremediation, biocatalysis, biofuel production, protein production, and biomedical research (Cubillos, 2016; Johnson & Echavarri-Erasun, 2011; Turker, M. 2015).

Yeast diversity influences the properties of fermented foods and beverages. For example, in wine production, *Saccharomyces cerevisiae* is responsible for alcoholic fermentation of grape, *Pichia* yeasts produce metabolites that add desirable organoleptic properties, while *Brettanomyces bruxellensis* is associated with unwanted off flavors (Burini et al., 2021; Ciani & Comitini, 2011; Gallone et al., 2016; Godoy et al., 2017; Jolly et al., 2014; Molinet & Cubillos, 2020). Moreover, yeast populations associated with human-related environments show specific adaptations. For instance, there are *S. cerevisiae* wine lineages that are tolerant to the presence of copper, sulfates, or high ethanol concentrations, while others used for beer production have the capacity to metabolize uncommon sugars such as maltose or maltotriose (Cubillos et al., 2019; De Chiara et al., 2022; Gallone et al., 2016; Giannakou et al., 2020, 2021; Liti et al., 2009; Peter et al., 2018; Warringer et al., 2011). Yeast domestication and its mechanisms have been most thoroughly documented in extreme scenarios of human handling, such as the use of starter cultures of *S. cerevisiae* in the production of beer (Gallone et al., 2016; Molinet & Cubillos, 2020).

In Mexico, agave spirits are obtained from the fermentation of juice and bagasse from cooked hearts of agave plants (*Agave* spp., Asparagaceae). This refers to all distilled beverages obtained from agave, such as *tequila*, *mezcal*, *bacanora*, *raicilla*, and other locally named spirits. Traditional production takes place in small or medium-sized distilleries across the country, where open fermentation occurs “spontaneously” by microorganisms from the surroundings living in a wide diversity of climates. An extensive range of production techniques are employed (Arellano-Plaza et al., 2022; CONABIO, 2006) and over fifty different agave taxa are used (Colunga-GarcíaMarín et al., 2017). In brief, agave spirit production involves plant harvesting, cooking, milling, fermentation, and distillation (Kirchmayr et al., 2017; Nolasco-Cancino et al., 2018). Cooking transforms complex carbohydrates such as fructans into simple sugars, mainly fructose. This step also gives rise to inhibitory compounds for microorganisms such as furans, aldehydes, and organic acids (Mancilla-Margalli & López, 2002). Through milling, pulp, fibers, and juice are obtained, and with the addition of water, open fermentation can proceed by the activity of environmental yeasts and other microorganisms (Kirchmayr et al., 2017; Lappe-Oliveras et al., 2008; Nolasco-Cancino et al., 2018; Páez-Lerma et al., 2013).

Despite an increasing demand for agave spirits in international markets, the diversity of yeasts responsible for agave fermentation remains largely unexplored. Pioneering studies by Marc-André Lachance in a traditional *tequila* distillery showed that local plants, insects, and objects in the distillery can work as the reservoirs of yeasts in agave fermentation (Lachance, 1995). In following efforts, yeasts have been isolated from fermentation tanks in some traditional distilleries in the states of Oaxaca, Durango, and Tamaulipas (Kirchmayr et al., 2017; Lappe-Oliveras et al., 2008; Nolasco-Cancino et al., 2018; Páez-Lerma et al., 2013). From these studies, at least 43 yeast and over 50 bacterial species have been associated with agave fermentations. Among these, *S. cerevisiae*, *Kluyveromyces marxianus* and *Pichia* species are the most abundant, regardless of the location sampled or year of the study.

The YeastGenomesMx consortium aims to provide a broad description and deeper understanding of the genomic and functional diversity of yeasts in Mexico. Considering the significance of these microorganisms in agave fermentations and the limited information available about their natural history in such environments, these communities are a good starting point to investigate yeast diversity in the region. For this purpose, we assembled the YMX.1.0 culture collection, comprising thousands of yeast strains isolated from agave fermentation tanks at 68 traditional distilleries across Mexico. Identification by matrix assisted laser desorption ionization time-of-flight (MALDI-TOF) mass spectrometry followed by ITS and D1/D2 sequencing of a subset of the isolates confirmed the regular occurrence of *S. cerevisiae* and revealed widespread presence of non-conventional yeasts, including species not previously associated with the agave environment and two candidate new species. The YMX.1.0 culture collection provides a unique resource for launching genomic and population studies of yeasts from agave fermentations in the context of a megadiverse region. As a working example, we generated and analyzed draft genome sequences of four isolates of the non-conventional yeast *Kazachstania humilis*, which showed that they are markedly different from strains from other regions.

## MATERIALS AND METHODS

### Agave distilleries and site information

Sampling sites were selected to include representative traditional distilleries from each of the seven producing regions of agave spirits in Mexico according to Aguirre-Dugua et al. (2006), with the West area subdivided (West I and West II) because of contrasting environmental conditions and agave species employed. Field work took place between March 2018 and December 2020. Samples were from 68 distilleries distributed across 13 states of Mexico (Colima, Durango, Estado de México, Guanajuato, Guerrero, Jalisco, Michoacán, Oaxaca, Puebla, San Luis Potosí, Sonora, Tamaulipas, and Zacatecas). Biogeographical parameters were obtained by sampling the geographical coordinates of the site over the corresponding raster cartography. Elevation data was obtained at 1 km scale resolution through the US Geological Survey GTOPO30 dataset (earthexplorer.usgs.gov; entities GT30W100N40 and GT30W140N40). Environmental data was obtained through WorldClim climatic layers (worldclim.org/bioclim) at 1 km (30 arc-seconds) resolution. Variables analyzed include Mean Annual Precipitation (wcbio_12), Mean Annual Temperature (wcbio_01), and Isothermality (wcbio_03); climates were classified as Aw (warm subhumid), (A)C (semiwarm), C(w) (temperate), and BS (arid and semiarid). The metadata of the location associated with each isolate is provided in **Dataset S1**.

### Collection and handling of agave fermentation samples

Samples were collected from distilleries with at least one active fermentation tank at the time of the visit. When more than one active tank was available, the agave species and fermentation phase information as provided by the producers was used to select the tanks to be sampled, namely one of each species or stage were sampled. The total number of sampled tanks was 181, ranging from one to five per distillery. Sampling was conducted by collecting agave must at arm’s length to the edge of the tank and 25-30 cm depth from the surface, using a sterile 25 mL serological pipette. Collected agave must was used to fill two prelabeled 4 mL cryovials with glycerol at a final concentration of 12.5%, which were immediately frozen in liquid nitrogen. All samples were kept in liquid nitrogen until they were transferred to a −80 °C freezer in the laboratory. In addition, site and sample data were collected in the field, including geographical coordinates, sample temperature, and pH (see **Dataset S1**).

### Yeast isolation and generation of the YMX.1.0 culture collection

Samples were thawed and 500 μL were used to inoculate Saccharomyces isolation medium. We followed a previously described isolation protocol (Liti et al., 2017) but lowered the concentration of ethanol in the medium to 6% v/v. A total of 158 out of 201 samples (86.7%) from 147 different tanks showed signs of fermentation after two to up to seven days; those samples were diluted 1:100 and 1:1000 and spread in Wallerstein Laboratory (WL) Nutrient Agar (Sigma 17222) which enables screening for morphological and physiological variation (Verdugo Valdez et al., 2011). Samples were incubated for 5 days at 25 °C and 2 days at 4 °C to increase morphological differentiation. Colonies showing different morphology were randomly selected from the plate showing the largest number of isolated colonies; 12-60 colonies were transferred to 150 μL of YPD medium with 12.5% glycerol in 96 microtiter-well plates (Corning CLS3367) and stored at −80°C. The distribution of the number of isolates per sample was 12, 24, 36 (43), 48, and 60 isolates for 4, 101, 43, 8, and 2 samples, respectively. We note that only those strains used for DNA extraction and ITS or D1/D2 sequencing were restreaked twice for isolated colonies. The YMX.1.0 culture collection contains 4,524 strains arranged in fifty-three microtiter plates.

### Strain identification by MALDI-TOF mass spectrometry

All strains were analyzed as in (García-Gamboa et al., 2021). In brief, microtiter plates were thawed, strain arrays were printed onto YPD agar plates and incubated for 48 h at 30 °C. Biomass was spotted directly onto MSP 96 target polished steel MALDI target plates; 1 μL of formic acid at 70% was added to each spot until completely dry, then spots were covered with 1 μL of 10mg/mL HCCA (α-cyano-4-hydroxycinnamic acid) matrix solution, 50% acetonitrile and 2.5% trifluoroacetic acid as organic solvent. Plates were processed in a MICROFLEX LT instrument (Bruker Daltonics) and biomass fingerprints were acquired for each spot in flexControl 3.4 software with the MBT_FC.par method. Forty laser shots were collected at six random positions within the spot, automatically generating mass spectra in the mass range from 2,000 to 20,000 Da. Output files were compared to the BDAL v.8 or v.10 databases (see **Dataset S1**). No mass peaks were obtained for 462 (10.2%) samples; for the remaining 4,524 samples, 3,063 (67.7%) passed a stringent Score Value of 1.7 that was used to assign a species for the isolate. A lenient Score Value of 1.5 resulted in 3,623 (80.0%) strain identifications.

### Strain identification by ITS and D1/D2 sequencing

A panel of 474 strains including four of the most abundant putative species identified by MALDI-TOF was selected. Isolates were restreaked two times to obtain axenic cultures, from which genomic DNA was extracted with sodium hydroxide (Sylvester et al., 2015). The ITS/5.8 rDNA ITS region was amplified using universal fungal primers ITS-1 and ITS-4, as previously described (Sylvester et al., 2015), and sequenced using the same primers. Sequences were filtered and processed using a 0.01 trimming threshold, using a modified Mott (M1) cut algorithm implemented in the *sangeranalyseR* package for R (Chao et al., 2021). For a subset of isolates that yielded no reliable identification by MALDI-TOF, taxonomic identity was assigned by analysis of the sequences of the ITS-5.8 S region as mentioned above and the D1/D2 variable domains that were amplified and sequenced using the NL1 and NL4 primers. The sequences were assembled, edited, and aligned with MEGA11 (Tamura et al., 2021) and compared with annotated yeast sequences in the GenBank database using BLAST, as previously described (Sayers et al., 2020). A low-depth sequencing approach was used for the taxonomic assignment of the isolates YMX003144, YMX000711, and YMX000142, which were assigned to two candidate new species. Briefly, genomic DNA was purified using the MasterPure DNA purification kit as recommended by the manufacturer and sequenced using the IlluminaNextSeq platform (2 x 150 bp). Poor quality reads were removed with fastp v0.20.0 (Chen et al., 2018). The remaining reads were assembled with SPAdes v3.12.0 (Bankevich et al., 2012). Nucleotide sequences of the ITS and D1/D2 regions were recovered from the assemblies with biostrings package using the sequence of the corresponding primers.

### Genome sequencing and analysis

Genome sequencing of three *Kz. humilis* strains was performed using Illumina and BGI platforms (2×150 pb). Raw reads were filtered by quality using fastp v0.20.0 (Chen et al., 2018). De novo assemblies of the filtered reads were performed using SPAdes v3.12.0 (Bankevich et al., 2012) with the default parameters. The preliminary assemblies were further scaffolded with RagTag v2.1.0 (Alonge et al., 2022) using the YMX004033 genome assembly (García-Ortega et al., 2022) as a reference. The assembled genomes, including the reference genomes (CLIB 1323 and YMX004033), were annotated using the MAKER pipeline v2.31.8 (Cantarel et al., 2008). The quast program was used to calculate assembly statistics. Pairwise ANI values between the strains and the type strain (CBS 1323) were determined using FastANI (Jain et al., 2018) and plotted with pheatmap packages (Kolde, 2012). Nucleotide sequences of the ITS, D1/D2 LSU rRNA, RPB1, RPB2, and EF-1α genes were downloaded from GenBank. When the sequences were not available, we used the biostrings R package to retrieve them from the whole-genome sequences. Alignments for the phylogenetic tree were performed with Clustal(Larkin et al., 2007). These alignments were concatenated and positions with gaps removed. The resulting alignment was used to infer a Maximum Likelihood tree with RAxML (Stamatakis, 2014). The GTR+GAMMA model for nucleotide substitution and bootstrap support values of 1,000 replicates was used. Structural variation between the CLIB1323 and YMX004033 genomes was identified through comparisons of whole-genome assemblies of both genomes using the SyRI software (Goel et al., 2019). We used ragtag to align and classify the scaffolds of each assembly and join them to produce comparable assemblies. Any unclassified scaffolds were eliminated from the synteny analysis, resulting in a comparable set of 13 scaffolds for each strain.

## RESULTS

### A comprehensive collection of yeast strains isolated from agave fermentations

The YMX.1.0 culture collection contains 4,524 yeast strains from sixty-eight artisanal agave distilleries in Mexico. Each isolate is linked to metadata regarding its preliminary species-level identification, biogeographical information, physicochemical parameters of the fermentation sample, and production practices (**Dataset S1**). Field work took place between 2018-2020 and a detailed description is provided in the Methods section.

Sampling sites were traditional distilleries distributed in twelve states across the seven regions in which agave spirits are produced in Mexico, namely the Northwest, Northeast, West I, West II, Balsas-basin, Central-highlands, and South-central regions (**Figure 1A**). *Tequila* distilleries were not considered given that, nowadays, axenic monocultures are typically used for controlled fermentation in the highly industrialized settings of these factories. The longest distance between sampling sites was 2,033 km (1,263 mi) from Baviácora, Sonora to San Luis Amatlán, Oaxaca. At least four general climate types were represented: tropical, subtropical, semiarid, and temperate. Sampling sites had an environmental annual temperature range between 14 and 24 °C with isothermality ranging from 49 to 78. The average annual rainfall was from 389 L/m^2^ in San Luis de la Paz, Guanajuato (Central highlands) to 1,510 L/m^2^ in Cabo Corrientes, Jalisco (West II), while altitudes were between 395 MASL in San Pedro Ures, Sonora (Northwest) to 2,508 MASL in San Felipe, Guanajuato (Central highlands) (**Figure 1B**). The diversity of production practices is illustrated by the types of fermentation tanks, which were built from wood, cement, plastic, stone, clay, steel, or cowhide (**Figure 1C**). Agave fermentation samples were nitrogen-frozen with glycerol in the field and used in the laboratory to isolate yeast strains (see Methods) (**Figure 1E**).

**Figure 1.**
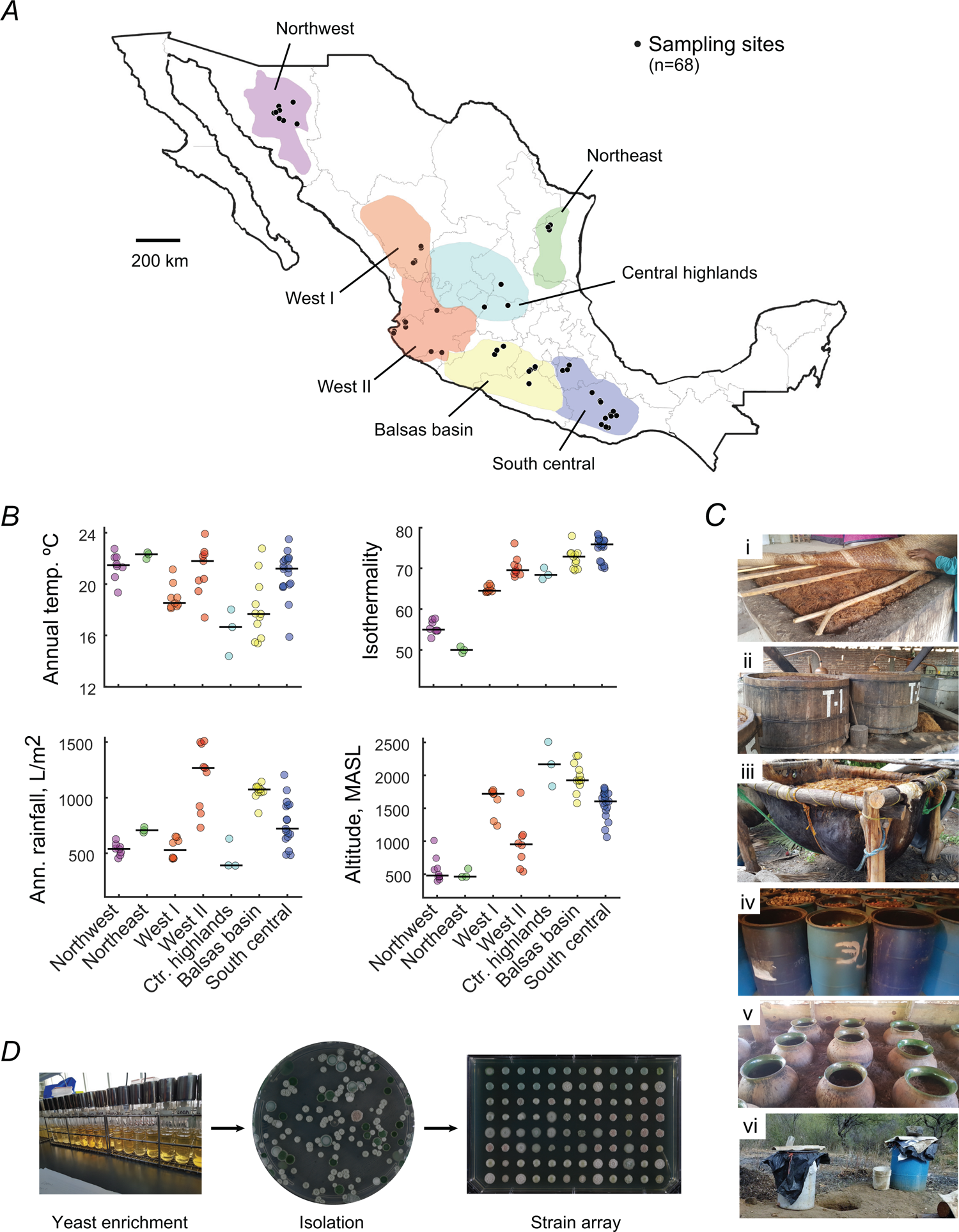
The YMX.1.0 culture collection was generated with yeasts isolated from diverse environmental and production conditions. (*A*) Sampling sites were 68 traditional agave distilleries across Mexico (black dots); shaded colored areas indicate the seven regions where agave spirits are produced in the country. **(*B*)** Scatter plots showing the diversity of climatic parameters of sampling sites grouped by region: Average annual temperature (top left), isothermality (top right), average annual rainfall (bottom right), and altitude above sea level (bottom left). **(*C*)** Samples were collected from actively fermenting tanks; pictures show examples of containers including (i) cement tanks, (ii) wood barrels, (iii) cowhide containers, (iv) plastic barrels, (v) clay pots, (vi) and steel barrels. **(*D*)** Fermented-agave samples were enriched in liquid growth medium and plated onto WL Nutrient Agar; isolated colonies were grown and stored in microtiter 96-well plates (see Methods).

### Identification and distribution of yeasts isolated from agave fermentation

To provide a comprehensive taxonomic description of the YMX.1.0 collection, we sought to characterize all isolated strains using MALDI-TOF mass spectrometry (**Dataset S1**). A minor number of samples (465; 10.3%) yielded no mass-spectrometry peaks and 996 (22.0%) samples provided spectra with no reliable identification in the database (Score Value cutoff <1.7). Importantly, 3,063 (67.7%) strains were classified using this stringent cutoff, corresponding to eighteen different yeast species. The collection also contained a small fraction of bacterial isolates (*n*=39). The number of classified isolates increased to 3,503 (77.4%) when using a 1.5 lenient cutoff, belonging to twenty-three yeast species. From the stringent set of classified strains, the most abundant putative species were *S. cerevisiae* (55.4 %) *Pichia kudriavzevii* (14.2 %), *Kl. marxianus* (8.6 %) and *Pichia manshurica* (6.6%) (**Figure 2A**; **Table 1**). In addition, at least 1% of the strains in the collection were assigned to *Pichia kluyveri*, *Torulaspora delbrueckii*, *Hanseniaspora lachancei*, *Pichia membranifaciens*, and *Hanseniaspora valbyensis*.

**Figure 2.**
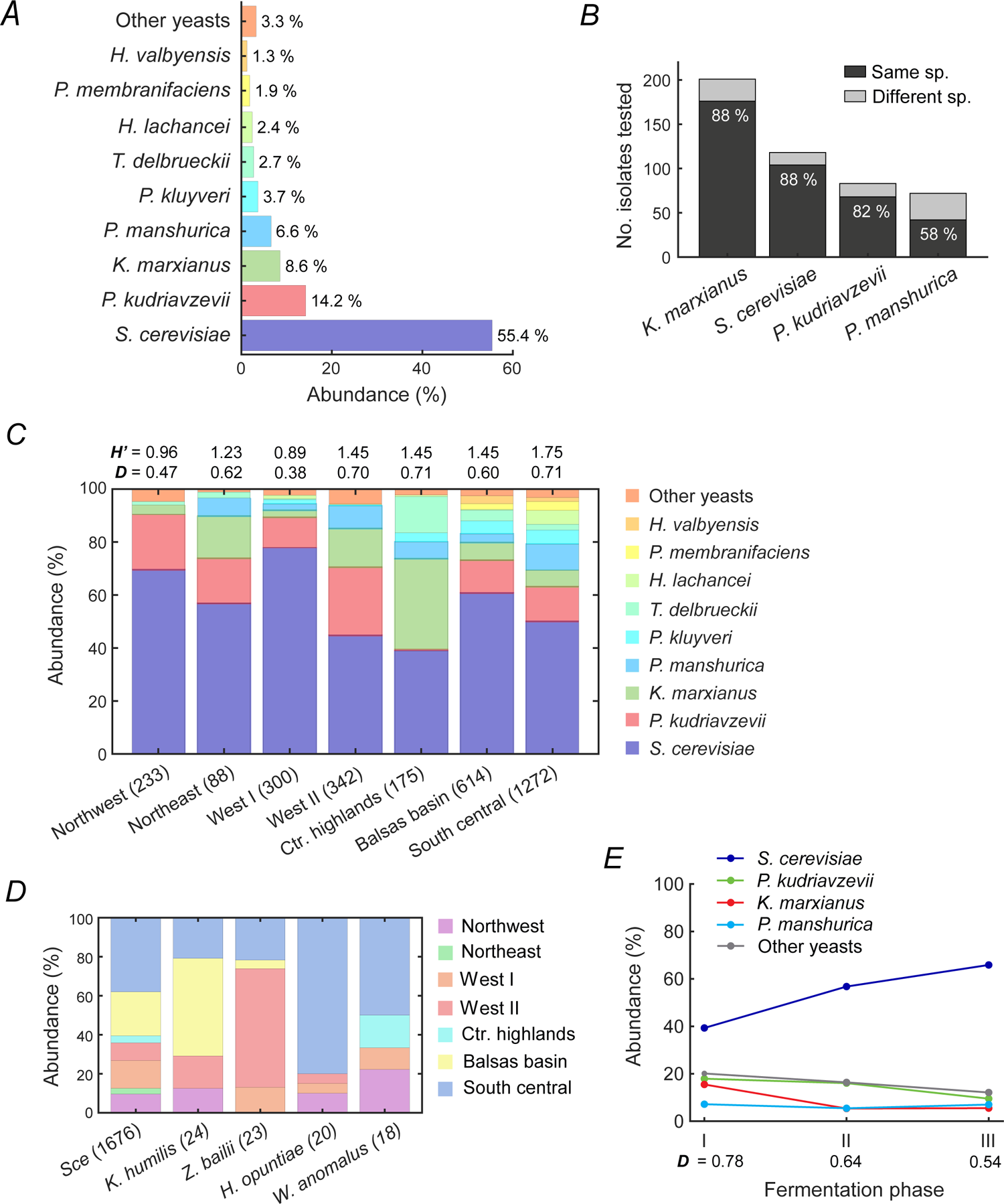
Species composition remains largely constant across geographic regions and fermentation phases. Using MALDI-TOF biotyping, 3,063 isolates of the YMX.1.0 culture collection were assigned to putative species. **(*A*)** Bar plot shows the frequency of species representing at least 1% of the isolates; other species comprised 3.3% of the identified strains. **(*B*)** A subset of 474 isolates assigned to the most abundant species were confirmed by ITS sequencing; plot shows the frequency of true (dark gray) and false (light gray) classifications. Correct species based on ITS sequencing are provided in Table S1. **(*C*)** Stacked bars show the species distributions of isolated yeasts in each of the seven producing regions; the number of isolates from each region are shown in parenthesis. The Shannon-Weiner (*H’*) and Simpson (*D’*) diversity indexes are indicated for each region (top). **(*D*)** Stacked bars show the frequencies of producing regions where the four species were isolated. These species were isolated at lower frequencies (0.5-1.0%) but were nonetheless present in at least 5% of the sampling sites; the number of isolates of each species are shown in parenthesis. *S. cerevisiae* data (*Sce*) is shown as a reference. **(*E*)** Plot shows the frequencies of the most abundant species at different phases of the fermentation; all other yeasts are grouped and plotted in a single data series (gray line). Three fermentation phases are defined from the ranked sampling times expressed as the fraction of total fermentation time in days.

**Table 1.** Saccharomycetales yeasts from the agave fermentation culture collection YMX.1.0 ^1^

| Species | Number of isolated strains | Frequency in distilleries (n=68) | Frequency in producing regions (n=7) |
| --- | --- | --- | --- |
| *Saccharomyces cerevisiae* * | 1,676 | 60 | 7 |
| *Pichia kudriavzevii* * | 430 | 41 | 7 |
| *Kluyveromyces marxianus* * | 259 | 30 | 7 |
| *Pichia manshurica* * | 199 | 33 | 6 |
| *Pichia kluyveri* | 111 | 21 | 5 |
| *Torulaspora delbrueckii* | 83 | 14 | 6 |
| *Hanseniaspora lachancei* | 73 | 8 | 2 |
| *Pichia membranifaciens* | 56 | 8 | 2 |
| *Hanseniaspora valbyensis* | 38 | 5 | 2 |
| *Kazachstania humilis* | 24 | 7 | 4 |
| *Zygosaccharomyces balii* | 23 | 5 | 4 |
| *Hanseniaspora opuntiae* | 20 | 6 | 4 |
| *Wickerhamomyces anomalus* | 18 | 6 | 4 |
| [*Candida*] *boidinii* / *Ogataea* clade | 5 | 2 | 2 |
| *Pichia fermentans* | 4 | 3 | 3 |
| *Pichia occidentalis* | 2 | 2 | 1 |
| *Wickerhamomyces subpelliculosus* | 2 | 2 | 1 |
| *Saccharomyces paradoxus* | 1 | 1 | 1 |

<sup>1</sup> Isolates were characterized by MALDI-TOF; data were obtained for 4,062 (89.8%) samples; species were assigned for 3,063 isolates (67.7%) using a stringent cutoff (Score Value>1.7)
\* A fraction of the isolates was assigned to a different species when analyzed by ITS sequencing (Table S1).

To control for species classification based on MALDI-TOF biotyping, we sequenced the rDNA internal transcribed spacers (ITS region) of a subset of 474 strains (**Table S1**). ITS sequences revealed that 390 (82.3%) of the isolates were correctly identified initially by MALDI-TOF biotyping at the species level (**Figure 2B**). The discrepancies between the methods are likely due to a combination of errors in sample handling and variations in the accuracy of the MALDI-TOF approach, which depends on the species, sample processing, and comprehensiveness of the reference database. The highest frequency of true identifications was for strains classified as *S. cerevisiae* (88.1%) and *Kl. marxianus* (87.6%), while the least accurate was *P. manshurica* (58.3%). Twenty-five out of the 72 putative *P. manshurica* strains (∼35%) corresponded to other *Pichia* species. Noteworthy, a small fraction (7.6%) of the putative *S. cerevisiae* were assigned to its sister species *Saccharomyces paradoxus* by ITS sequencing, increasing the total number of *S. paradoxus* isolates to ten in the YMX.1.0 collection.

Based on MALDI-TOF classification data, the most abundant yeast species were widely distributed in fermentation tanks located across the seven producing regions. Despite the great diversity of environmental conditions, the species composition of the yeast communities in distilleries across sampling regions remained largely constant (*p*=0.204, *r*=0.040; ANISOM) (**Figure 2C**; **Table 1**). Using the same data, we calculated the Shannon-Weiner (*H’*) and Simpson (*D’*) indexes for each region, showing that yeast species diversity varied from *H’*=0.96 in the Northwest to *H’*=1.75 in the South-central region. Dominance of the most abundant yeast species was marked in the Northwest (*D*=0.47) and less evident in other regions. Noteworthy, isolation of the otherwise widespread *P. kudriavzevii* yeast was uncommon in the Central-highlands region; likewise, *P. manshurica* was uncommon in the Northwest region. In contrast, *T. delbrueckii* and *H. lachanceii* isolates were common in the Central-highlands and South-central regions, respectively, although both species were overall rare. Other species were isolated at lower frequencies (0.5-1%) but were nonetheless distributed across different sampling sites and regions (**Figure 2D**; **Table 1**).

To test whether the fermentation phase influenced the yeast diversity of the isolated strains in the sampled tanks, we compared the diversity of isolates from different samples that were collected at different relative times during the fermentation process (**Figure 2E**). The Simpson diversity index decreased as a function of fermentation time, which could be explained in part by an increase in the isolation frequency of *S. cerevisiae* strains in late compared to early fermentations. It is important to note that these comparisons were made using samples collected from distinct locations, which could have influenced the observed trends. Nonetheless, the diversity changes were so subtle that it seems that there were no major changes in the frequencies of yeast species as a function of fermentation time, at least for the most abundant species. Although further investigation is needed, together, these results suggest there is a common core of yeast species in agave fermentations regardless of the fermentation phase and the environmental and biological diversity of the sites sampled.

### New species of yeasts in the agave fermentation environment

Many isolates of the YMX.1.0 with good quality spectra (996, 22.0%) were not assigned to any species in the database using a high confidence threshold (ScoreValue>=1.7). This number was still considerable when using a lenient cutoff of >=1.5 resulting in 439 isolates with unreliable identification. To explore the diversity of yeasts in this set of non-identified strains, the D1/D2 domains and ITS region of 22 isolates from Oaxaca (South-central region) and Durango (West I region) with no MALDI-TOF classification were sequenced. Analysis of the sequences showed that the strains isolated belong to three genera, namely *Saccharomyces*, *Torulaspora*, and *Pichia*. Six species were identified, four of which were previously known while two were candidates for novel species (**Table 2**).

**Table 2.**
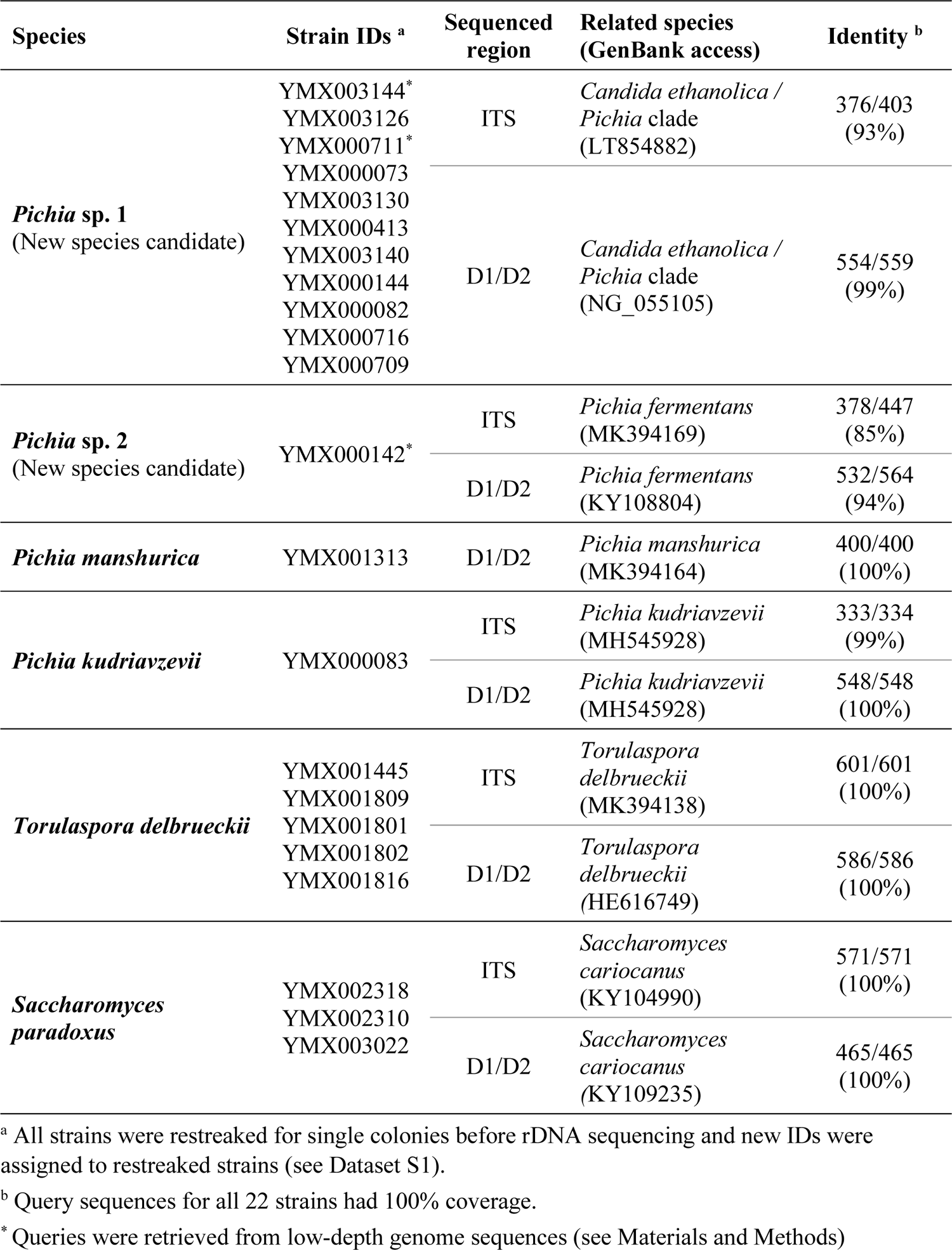
Identification of 22 Saccharomycetales isolates from agave fermentation based on sequences of the ITS region and D1/D2 domains of the large subunit of the rRNA gene

In this set of yeast isolates, three showed high identity to a strain isolated from fruit fly formerly classified as *S. cariocanus*; this species has been reclassified as *S. paradoxus* based on the low sequence divergence between the strains (Liti et al., 2006). In addition, five strains from the South-central region were identified as *T. delbrueckii*; this species has been previously reported in open agave fermentations (Muñoz-Miranda et al., 2022). Regarding the *Pichia* clade, three species were represented by a single isolate, namely *P. kudriavzevii*, *P. manshurica*, and a new species candidate *Pichia* sp. 2 YMX000142. We observed that the nearest relative of this putative new species was *Pichia* sp. 2 (YMX000142). We observed that the nearest relative of this putative new species was *P. fermentans*, from which it differed by 32 substitutions in the D1/D2 domains and 69 substitutions in the ITS region. Interestingly, query sequences of eleven isolates from the two producing regions were identical and represented an additional new species candidate, *Pichia* sp. 1, closely related to the [*Candida*] *ethanolica / Pichia* clade. The candidate new species *Pichia* sp. 1 (YMX003144 and others) differed by 27 substitutions in ITS sequences and 5 substitutions in D1/D2 domains from *C. ethanolica*. These two candidate new species isolated will be formally described elsewhere. Together, these results suggest that the sub-sampled agave fermentation environment harbors new-to-science species of fungi.

### Genome analysis reveals substantial intraspecific genetic diversity in *Kazachstania humilis*

The YMX.1.0 culture collection provides a unique biological resource for performing genomic and population studies of the different yeast species found in agave fermentations. The yeast *Kz. humilis* (*Candida humilis*, *Candida milleri*) is one of the dominant species in type I sourdoughs (Carbonetto et al., 2020; García-Ortega et al., 2022; Wittwer et al., 2022) and one of the most abundant fungi during the preparation of a variety of fermented vegetables and beverages in China (Guan et al., 2020; Gullo et al., 2003; Wittwer et al., 2022), but few occurrences of the species have been reported in the Americas. In our study, the species was isolated with low frequency (24 isolates in the complete collection) but was present in seven (10%) distilleries distributed in four of the seven producing regions (**Table 1**).

To gain a better understanding of the genomic diversity of *Kz. humilis* from agave fermentations, we conducted whole genome sequencing and analysis of four strains from distant regions, including the previously sequenced YMX004033 strain (García-Ortega et al., 2022). The obtained assemblies were compared with the genome sequence of the type-strain CLIB 1323 isolated from sourdough in France (Jacques et al., 2016). The draft genome sizes of the three new *Kz. humilis* isolates sequenced in this study (YMX00387, YMX000554 and YMX003162) varied from 17.1 to 19 Mbp, a larger size than that of the YMX004033 and CLIB 1323 genome references (13.9 Mbp) (**Table S2**). The variable sizes possibly reflect the degree of fragmentation of the genomes derived from this study (229 scaffolds in average), in comparison to those obtained with long-reads, YMX004033 (21 scaffolds) and CLIB 1323 (16 scaffolds). The average number of predicted genes in the four strains from agave-fermentation was 5,333, which is higher than the type strain’s predicted gene count of 4,295. However, this value is consistent with the number of genes reported for other species within the same genus, including *Kz. barnettii* (5,276), *Kz. exigua* (5,416), *Kz. unispora* (5,287), *Kz. saulgeensis* (5,329), *Kz. africana* (5,375), and *Kz. naganishii* (5,319).

To better understand the phylogenetic position of the four *Kz. humilis* strains from agave fermentation, we retrieved the nucleotide sequences of five commonly used genetic markers (ITS, D1/D2 LSU rRNA, RPB1, RPB2, and EF-1α) from the genome assemblies, concatenated them, and reconstructed a phylogenetic tree from their alignment with other available sequences of *Kazachstania* species (**Figure 3A**). The reconstructed phylogenetic tree was congruent with phylogenies previously reported for these species (Jacques et al., 2016). As expected, strains from agave fermentations grouped together in one clade, with the CLIB 1323 *Kz. humilis* type strain and CBS 5658 and NRRL Y-7245 isolates as sister taxa.

**Figure 3.**
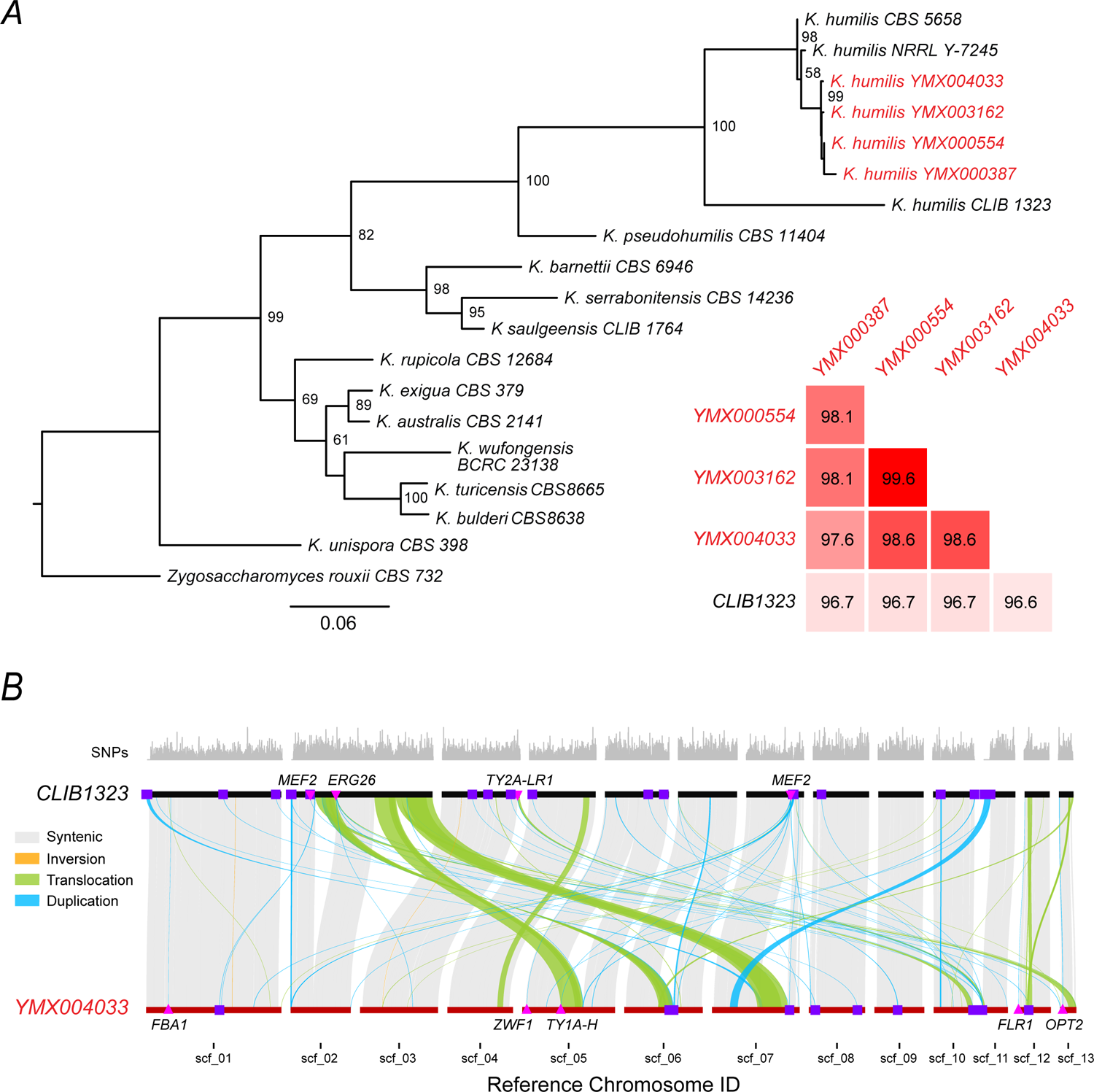
Comparative genomics of *Kazachstania humilis* isolates. (*A*) Maximum-likelihood phylogenetic tree for *Kazachstania* species based on the alignment of the concatenated ITS, D1/D2 LSU rRNA, RPB1, RPB2 and EF-1a genes (2,863 positions in total); *Zygosaccharomyces rouxii* CBS 732 was used as the outgroup. The four strains from agave fermentation are shown in red. Numbers on the branches are bootstrap support based on 1,000 replicates; only numbers higher than 50% are shown. The genetic-distance scale represents substitutions per nucleotide. The matrix (inset, bottom right) shows the genome-wide pairwise average nucleotide identity (ANI) values for *Kz. humilis* strains with available genome sequences. **(*B*)** Genome comparison between the genomes of the type strain *Kz. humilis* CLIB 1323 (black lines) and strain YMX004033 (red lines). Figure shows syntenic (gray), duplicated (cyan), translocated (green), and inverted (orange) regions larger than 10 Kb. Duplicated regions resulting in gene-copy gains are indicated by magenta triangles, while purple squares overlap duplicated regions with simple and low complexity repeat elements. Names of duplicated genes are shown next to the genome where the duplication is observed. SNP densities of CLIB 1323 are shown at the top, in non-overlapping 1 Kb bins.

To provide a quantitative measure of the genetic relationship between the *Kz. humilis* strains from agave and the type strain at the genome-wide level, we calculated the average nucleotide identity (ANI) between all strains with available genome sequences. The ANI values ranged from 96.5 to 99.56 % (**Figure 3A, inset**), which are values within the range of diversity expected for yeast strains that belong to the same species (Gostinčar, 2020). The highest ANI value was found for YMX000554 and YMX003162 (99.56 %), two agave yeast strains from the same producing region. The CLIB 1323 type strain had the lowest median ANI value with respect to all strains from agave fermentations (96.68 %), suggesting that the agave and sourdough strains could be from different populations of *Kz. humilis*.

To identify structural variation between the genomes of these strains, we performed a synteny analysis between the two strains with available genome sequences assembled from long-read data: the *Kz. humilis* type strain CLIB 1323 (Jacques et al., 2016) from sourdough in France and the agave-fermentation isolate YMX004033 from the West II producing region in Mexico (García-Ortega et al., 2022) (**Table S3**). Analysis of syntenic regions across 13 consistent scaffolds revealed ∼11.8 Mb of syntenic regions (∼85.9% of the CLIB 1323 genome), 66 translocations, 137 duplications in CLIB 1323 and 71 duplications in YMX004033, 57,063 INDELs and 50,3408 SNPs. In addition, SyRI identified 0.24 Mb of CLIB 1323 and 0.69 Mb of YMX004033 DNA sequences that were not aligned with the other respective genome (**Figure 3B**). These major structural variants further suggest that the *Kz. humilis* strains belong to different populations, with potential reduced genetic compatibility between them. Structural variation also included several segmental and dispersed duplications, some of them encompassing complete open reading frames. Together, these findings indicated that the strains isolated from agave fermentations are genetically distinct, underscoring the potential of the YMX.1.0 culture collection for future research on population genomics of non-conventional yeast species that are important for food and beverage production worldwide.

## DISCUSSION

The diversity of yeasts associated with agave fermentation has remained underexplored, despite the widespread biological and industrial interest in the Saccharomycetales yeasts and an increasing demand for agave spirits in the global markets. In this study, we have described the yeast species present in this human-related environment by systematically isolating and identifying thousands of strains from traditional distilleries across Mexico. Each isolate of the YMX.1.0 culture collection is linked to metadata for the fermentation sample and isolation site including biogeographical information, physicochemical parameters, and production practices. Despite the great diversity of environmental conditions, number of agave species used, and variety of production practices, the composition of yeast communities remained largely homogeneous throughout sampled locations. This suggests that there is a core consortium of yeast species associated with these fermentations whose composition is more strongly influenced by the common characteristics of the fermentation substrate rather than by the specific surrounding environment and production practices of each distillery. However, it is possible that intraspecific genetic differentiation exists across producing regions and there may also be phenotypes associated with local adaptations. Each of the identified species may harbor undescribed populations and gene pools, as herein shown for the non-conventional yeast *Kz. humilis*.

The description of the communities in this study is based on cultured, isolated strains and is biased by the enrichment conditions used. Nonetheless, the species were in good agreement with previous studies using alternative culture-enrichment approaches (Kirchmayr et al., 2017; Nolasco-Cancino et al., 2018). Despite the limitations of the cultured-based method to characterize the overall yeast composition, the systematic nature and extension of our sampling effort offers an unprecedented opportunity to contrast the diversity of yeast species isolated from agave fermentations from diverse biogeographical and cultural contexts. Future studies using metagenomics approaches will provide a more complete description of the core yeast community in these fermentations and may reveal unique features of the microbial communities of specific producing regions. The YMX1.0 culture collection also opens new research avenues to understand crucial aspects of yeast biology such as genomic and functional diversity within populations that would not be possible using approaches that are not based on cultured organisms. For instance, gene duplication of the *ZWF1* gene in a *Kz. humilis* strain isolated here from an agave fermentation could be associated with tolerance to furans in this environment, as increased expression of genes of the pentose-phosphate pathway has been shown to underlie tolerance to hydroxymethylfurfural in *S. cerevisiae* (Gencturk & Ulgen, 2022; Gorsich et al., 2006; Park et al., 2011). The availability of a comprehensive collection of strains isolated from agave fermentations will now allow experimental testing of this type of hypothesis.

In this survey, yeast strains were taxonomically classified mostly by MALDI-TOF biotyping using direct biomass transfer (García-Gamboa et al., 2021), which allows rapid identification directly from single colonies. This approach results in higher rates of incorrect species assignment compared to sequencing-based methods. We evaluated the extent of this problem by sequencing the ITS region of a subset of 474 isolates and showed that disparate species were assigned in ∼19% of the cases. Noteworthy, García-Gamboa et al. (2021) also used yeast extracts for MALDI-TOF classification, achieving 100% correct identification of *P. kluyveri* isolates while comparing to ITS sequencing. In our study, incorrect classification was strongly biased to one of the species tested (*P. manshurica*), underscoring that the reference databases used for mass-spectrometry biotyping are yet to be improved for less studied species. This is especially relevant for the characterization of agave fermentations in which *Picha* species have been shown to be common. ITS and D1/D2 sequencing also revealed the occurrence of *S. paradoxus* in agave fermentations. As opposed to its sister *S. cerevisiae*, this species is relatively less common in fermentative environments (Dashko et al., 2016). Importantly, our survey revealed two new species candidates related to the *Pichia* clade, one of which seemed to be broadly present in two of the producing regions surveyed. Further work will be needed to characterize these isolates. These results suggest that the isolates from the YMX.1.0 culture collection are useful for the description of new fungal species from a still poorly characterized fermentative environment.

The YMX.1.0 culture collection offers a powerful resource for population genomics of the Saccharomycetales yeasts. Analyzing the many strains of model species such as *S. cerevisiae* or *Kl. marxianus* will help to understand their evolution and natural history at the interface of natural and human-related environments. In fact, genome sequences of a handful of isolates of *S. cerevisiae* from agave fermentation have already shown that they are considerably different with genomic features such as widespread introgressions (Peter et al., 2018). In the case of *Kl. marxianus*, the genome of a strain isolated from agave juice has been shown to form a specific clade, far apart from all other sequenced strains of this species (Ortiz-Merino et al., 2018). The YMX.1.0 culture collection is also an untapped resource to study the diversity of non-conventional yeast species. Here, we observed that *Kz. humilis*, which is commonly found in sourdoughs around the world (Wittwer et al., 2022), also thrives in agave fermentations. The draft genome sequences of four of these agave fermentation strains suggested that they belong to a different population given their genetic distance to sourdough isolates. Overall, yeast strains in the YMX.1.0 culture collection provide the opportunity to compare the adaptation trajectories of distinct species that have been exposed to the same environment.

Most of the emphasis in the study and conservation of agave spirit production has been placed on the plants and production practices employed. However, to preserve this system integrally we still need to consider the microbial communities that are essential for open fermentation associated with small and medium-sized distilleries. Our study provides a unique resource to grant deeper knowledge of the genomic and functional diversity of yeasts associated with this exceptional fermentation system in one of the megadiverse regions of the world. The YMX.1.0 collection directly enables the conservation of the microorganisms that are essential to produce agave spirits, an activity of crucial commercial and cultural importance for rural communities in Mexico. Future research based on our work will contribute to a better understanding of not only how the microbial communities contribute to the production of agave spirits, but also to the biology and natural history of microorganisms living at the interphase of natural environments and traditional distilleries.

## DATA AVAILABILITY

ITS and D1/D2 sequences of yeast isolates corresponding to two candidate new species *Pichia* sp. 1 and *Pichia* sp. 2 are available at GenBank database under accession numbers OR034128, OR225219, and OR034113, OR034133. The three new *Kazachstania humilis* genome sequences have been deposited at NCBI under accession numbers JAIWYT000000000, JAIWYS000000000, and JAIWYU000000000.

## ACKNOWLEDGEMENTS

We are indebted to over one-hundred producers of agave spirits in the states of Colima, Durango, Estado de México, Guanajuato, Guerrero, Jalisco, Michoacán, Oaxaca, Puebla, San Luis Potosí, Sonora, Tamaulipas, and Zacatecas, Mexico, who kindly participated in this research by sharing their knowledge and providing access to samples. For help with field work and sampling, we thank Karina Abad, Maritza Álvarez, Edgar Ángeles, Graciela Ángeles, Vanessa Arellano, Margareta Boege, Aida Carbajal, Jorge Carbajal, Carolina Castañeda, Bertha Castillo Ortega, Jason Paul Cox, Jorge Cuamatzi, Jilberto Dávila, Aarón De Luna, Marco Chávez, Hendrik Giersiepen, Alejandro González Anaya, Mathieu Harcot, Gerardo Hernández, Alejandro Juárez, Claudio López, Xavier López Medellín, Artemiza Martínez, Alberto Martínez López, José Martínez Rodríguez, Rodrigo Medellín, Alejandro Olmedo, Anibal Reyes, Benito Salcedo, Ivan Sedeño, Nelly Selem, José Antonio Urban, Carina Uribe, and Karen Vaca. We thank Luis Aguilar, René Delgado, and Jair García Sotelo for technical assistance and Raúl Ortiz-Merino for critical reading of the manuscript. We are grateful to Iván Saldaña for invaluable suggestions for field work and pilot sampling in Oaxaca and to Anne Gschaedler for her generous advice during the initial phases of the project.

## SUPPORTING INFORMATION

**Table S1.** Confirmation of species assignment by ITS sequencing (PDF file)

**Table S2.** Assembly statistics of four *Kazachstania humilis* strains from agave fermentation and the CLIB 1323 type strain (PDF file)

**Table S3.** Summary of SyRI structural annotation of *Kazachstania humilis* genomes (PDF file)

**Dataset S1.** The YMX.1.0 culture collection and associated metadata (XLSX file)

