## Supplementary material for "Yeast diversity in open agave fermentations across Mexico": Table S

**Table S1. Confirmation of species assignment by ITS sequencing**

| Strain ID | Score_MALDITOF | Species_MALDITOF | Species_ITS |
|---|---|---|---|
| YMX001636 | 1.71 | Kluyveromyces marxianus | Hanseniaspora lachancei |
| YMX002002 | 2.39 | Kluyveromyces marxianus | Kluyveromyces marxianus |
| YMX001896 | 2.322 | Kluyveromyces marxianus | Kluyveromyces marxianus |
| YMX001507 | 2.294 | Kluyveromyces marxianus | Kluyveromyces marxianus |
| YMX002752 | 2.281 | Kluyveromyces marxianus | Kluyveromyces marxianus |
| YMX002184 | 2.28 | Kluyveromyces marxianus | Kluyveromyces marxianus |
| YMX002182 | 2.271 | Kluyveromyces marxianus | Kluyveromyces marxianus |
| YMX002742 | 2.263 | Kluyveromyces marxianus | Kluyveromyces marxianus |
| YMX001888 | 2.26 | Kluyveromyces marxianus | Kluyveromyces marxianus |
| YMX002741 | 2.26 | Kluyveromyces marxianus | Kluyveromyces marxianus |
| YMX002352 | 2.254 | Kluyveromyces marxianus | Kluyveromyces marxianus |
| YMX002739 | 2.251 | Kluyveromyces marxianus | Kluyveromyces marxianus |
| YMX001062 | 2.243 | Kluyveromyces marxianus | Kluyveromyces marxianus |
| YMX002347 | 2.24 | Kluyveromyces marxianus | Kluyveromyces marxianus |
| YMX003030 | 2.24 | Kluyveromyces marxianus | Kluyveromyces marxianus |
| YMX001887 | 2.23 | Kluyveromyces marxianus | Kluyveromyces marxianus |
| YMX002348 | 2.222 | Kluyveromyces marxianus | Kluyveromyces marxianus |
| YMX001884 | 2.22 | Kluyveromyces marxianus | Kluyveromyces marxianus |
| YMX003033 | 2.22 | Kluyveromyces marxianus | Kluyveromyces marxianus |
| YMX002166 | 2.211 | Kluyveromyces marxianus | Kluyveromyces marxianus |
| YMX003031 | 2.21 | Kluyveromyces marxianus | Kluyveromyces marxianus |
| YMX003874 | 2.202 | Kluyveromyces marxianus | Kluyveromyces marxianus |
| YMX001592 | 2.2 | Kluyveromyces marxianus | Kluyveromyces marxianus |
| YMX003026 | 2.2 | Kluyveromyces marxianus | Kluyveromyces marxianus |
| YMX003036 | 2.2 | Kluyveromyces marxianus | Kluyveromyces marxianus |
| YMX003254 | 2.2 | Kluyveromyces marxianus | Kluyveromyces marxianus |
| YMX000772 | 2.19 | Kluyveromyces marxianus | Kluyveromyces marxianus |
| YMX002345 | 2.19 | Kluyveromyces marxianus | Kluyveromyces marxianus |
| YMX002390 | 2.19 | Kluyveromyces marxianus | Kluyveromyces marxianus |
| YMX001590 | 2.181 | Kluyveromyces marxianus | Kluyveromyces marxianus |
| YMX001078 | 2.18 | Kluyveromyces marxianus | Kluyveromyces marxianus |
| YMX001885 | 2.18 | Kluyveromyces marxianus | Kluyveromyces marxianus |
| YMX002142 | 2.18 | Kluyveromyces marxianus | Kluyveromyces marxianus |
| YMX003041 | 2.17 | Kluyveromyces marxianus | Kluyveromyces marxianus |
| YMX001065 | 2.164 | Kluyveromyces marxianus | Kluyveromyces marxianus |
| YMX001508 | 2.16 | Kluyveromyces marxianus | Kluyveromyces marxianus |
| YMX003029 | 2.16 | Kluyveromyces marxianus | Kluyveromyces marxianus |
| YMX001690 | 2.15 | Kluyveromyces marxianus | Kluyveromyces marxianus |
| YMX002754 | 2.15 | Kluyveromyces marxianus | Kluyveromyces marxianus |
| YMX003259 | 2.15 | Kluyveromyces marxianus | Kluyveromyces marxianus |
| YMX001686 | 2.14 | Kluyveromyces marxianus | Kluyveromyces marxianus |
| YMX003256 | 2.14 | Kluyveromyces marxianus | Kluyveromyces marxianus |
| YMX002170 | 2.132 | Kluyveromyces marxianus | Kluyveromyces marxianus |
| YMX001688 | 2.13 | Kluyveromyces marxianus | Kluyveromyces marxianus |
| YMX002418 | 2.124 | Kluyveromyces marxianus | Kluyveromyces marxianus |

| | | | |
|---|---|---|---|
| YMX003172 | 2.122 | Kluyveromyces marxianus | Kluyveromyces marxianus |
| YMX003045 | 2.12 | Kluyveromyces marxianus | Kluyveromyces marxianus |
| YMX002144 | 2.1 | Kluyveromyces marxianus | Kluyveromyces marxianus |
| YMX002329 | 2.1 | Kluyveromyces marxianus | Kluyveromyces marxianus |
| YMX003044 | 2.1 | Kluyveromyces marxianus | Kluyveromyces marxianus |
| YMX003094 | 2.1 | Kluyveromyces marxianus | Kluyveromyces marxianus |
| YMX002338 | 2.09 | Kluyveromyces marxianus | Kluyveromyces marxianus |
| YMX002138 | 2.08 | Kluyveromyces marxianus | Kluyveromyces marxianus |
| YMX002337 | 2.08 | Kluyveromyces marxianus | Kluyveromyces marxianus |
| YMX003257 | 2.073 | Kluyveromyces marxianus | Kluyveromyces marxianus |
| YMX002334 | 2.071 | Kluyveromyces marxianus | Kluyveromyces marxianus |
| YMX001691 | 2.07 | Kluyveromyces marxianus | Kluyveromyces marxianus |
| YMX003038 | 2.07 | Kluyveromyces marxianus | Kluyveromyces marxianus |
| YMX003241 | 2.06 | Kluyveromyces marxianus | Kluyveromyces marxianus |
| YMX001562 | 2.05 | Kluyveromyces marxianus | Kluyveromyces marxianus |
| YMX002150 | 2.05 | Kluyveromyces marxianus | Kluyveromyces marxianus |
| YMX002151 | 2.05 | Kluyveromyces marxianus | Kluyveromyces marxianus |
| YMX002152 | 2.05 | Kluyveromyces marxianus | Kluyveromyces marxianus |
| YMX002932 | 2.05 | Kluyveromyces marxianus | Kluyveromyces marxianus |
| YMX003255 | 2.05 | Kluyveromyces marxianus | Kluyveromyces marxianus |
| YMX002158 | 2.044 | Kluyveromyces marxianus | Kluyveromyces marxianus |
| YMX003185 | 2.044 | Kluyveromyces marxianus | Kluyveromyces marxianus |
| YMX001683 | 2.04 | Kluyveromyces marxianus | Kluyveromyces marxianus |
| YMX002147 | 2.034 | Kluyveromyces marxianus | Kluyveromyces marxianus |
| YMX003701 | 2.021 | Kluyveromyces marxianus | Kluyveromyces marxianus |
| YMX001579 | 2.02 | Kluyveromyces marxianus | Kluyveromyces marxianus |
| YMX001687 | 2.02 | Kluyveromyces marxianus | Kluyveromyces marxianus |
| YMX003034 | 2.02 | Kluyveromyces marxianus | Kluyveromyces marxianus |
| YMX003756 | 2.014 | Kluyveromyces marxianus | Kluyveromyces marxianus |
| YMX002156 | 2.01 | Kluyveromyces marxianus | Kluyveromyces marxianus |
| YMX003258 | 2.01 | Kluyveromyces marxianus | Kluyveromyces marxianus |
| YMX003841 | 2.01 | Kluyveromyces marxianus | Kluyveromyces marxianus |
| YMX004034 | 2.01 | Kluyveromyces marxianus | Kluyveromyces marxianus |
| YMX000998 | 2 | Kluyveromyces marxianus | Kluyveromyces marxianus |
| YMX001074 | 2 | Kluyveromyces marxianus | Kluyveromyces marxianus |
| YMX001584 | 2 | Kluyveromyces marxianus | Kluyveromyces marxianus |
| YMX001892 | 2 | Kluyveromyces marxianus | Kluyveromyces marxianus |
| YMX003048 | 2 | Kluyveromyces marxianus | Kluyveromyces marxianus |
| YMX002447 | 1.992 | Kluyveromyces marxianus | Kluyveromyces marxianus |
| YMX003848 | 1.992 | Kluyveromyces marxianus | Kluyveromyces marxianus |
| YMX001076 | 1.99 | Kluyveromyces marxianus | Kluyveromyces marxianus |
| YMX002336 | 1.99 | Kluyveromyces marxianus | Kluyveromyces marxianus |
| YMX003863 | 1.99 | Kluyveromyces marxianus | Kluyveromyces marxianus |
| YMX002143 | 1.98 | Kluyveromyces marxianus | Kluyveromyces marxianus |
| YMX002849 | 1.98 | Kluyveromyces marxianus | Kluyveromyces marxianus |
| YMX003251 | 1.98 | Kluyveromyces marxianus | Kluyveromyces marxianus |
| YMX004047 | 1.974 | Kluyveromyces marxianus | Kluyveromyces marxianus |

| | | | |
|---|---|---|---|
| YMX003243 | 1.972 | Kluyveromyces marxianus | Kluyveromyces marxianus |
| YMX001698 | 1.971 | Kluyveromyces marxianus | Kluyveromyces marxianus |
| YMX003248 | 1.97 | Kluyveromyces marxianus | Kluyveromyces marxianus |
| YMX003458 | 1.97 | Kluyveromyces marxianus | Kluyveromyces marxianus |
| YMX002141 | 1.964 | Kluyveromyces marxianus | Kluyveromyces marxianus |
| YMX002341 | 1.963 | Kluyveromyces marxianus | Kluyveromyces marxianus |
| YMX001201 | 1.96 | Kluyveromyces marxianus | Kluyveromyces marxianus |
| YMX003754 | 1.951 | Kluyveromyces marxianus | Kluyveromyces marxianus |
| YMX001267 | 1.95 | Kluyveromyces marxianus | Kluyveromyces marxianus |
| YMX003032 | 1.95 | Kluyveromyces marxianus | Kluyveromyces marxianus |
| YMX003247 | 1.95 | Kluyveromyces marxianus | Kluyveromyces marxianus |
| YMX004060 | 1.95 | Kluyveromyces marxianus | Kluyveromyces marxianus |
| YMX002139 | 1.94 | Kluyveromyces marxianus | Kluyveromyces marxianus |
| YMX001072 | 1.934 | Kluyveromyces marxianus | Kluyveromyces marxianus |
| YMX001252 | 1.93 | Kluyveromyces marxianus | Kluyveromyces marxianus |
| YMX002929 | 1.93 | Kluyveromyces marxianus | Kluyveromyces marxianus |
| YMX003667 | 1.93 | Kluyveromyces marxianus | Kluyveromyces marxianus |
| YMX004039 | 1.924 | Kluyveromyces marxianus | Kluyveromyces marxianus |
| YMX001573 | 1.923 | Kluyveromyces marxianus | Kluyveromyces marxianus |
| YMX002381 | 1.921 | Kluyveromyces marxianus | Kluyveromyces marxianus |
| YMX002200 | 1.92 | Kluyveromyces marxianus | Kluyveromyces marxianus |
| YMX003043 | 1.92 | Kluyveromyces marxianus | Kluyveromyces marxianus |
| YMX003788 | 1.915 | Kluyveromyces marxianus | Kluyveromyces marxianus |
| YMX003246 | 1.91 | Kluyveromyces marxianus | Kluyveromyces marxianus |
| YMX004049 | 1.903 | Kluyveromyces marxianus | Kluyveromyces marxianus |
| YMX001994 | 1.901 | Kluyveromyces marxianus | Kluyveromyces marxianus |
| YMX001648 | 1.9 | Kluyveromyces marxianus | Kluyveromyces marxianus |
| YMX002936 | 1.893 | Kluyveromyces marxianus | Kluyveromyces marxianus |
| YMX003208 | 1.89 | Kluyveromyces marxianus | Kluyveromyces marxianus |
| YMX002474 | 1.88 | Kluyveromyces marxianus | Kluyveromyces marxianus |
| YMX002937 | 1.88 | Kluyveromyces marxianus | Kluyveromyces marxianus |
| YMX003028 | 1.88 | Kluyveromyces marxianus | Kluyveromyces marxianus |
| YMX003790 | 1.88 | Kluyveromyces marxianus | Kluyveromyces marxianus |
| YMX003245 | 1.873 | Kluyveromyces marxianus | Kluyveromyces marxianus |
| YMX002330 | 1.87 | Kluyveromyces marxianus | Kluyveromyces marxianus |
| YMX002948 | 1.86 | Kluyveromyces marxianus | Kluyveromyces marxianus |
| YMX001891 | 1.85 | Kluyveromyces marxianus | Kluyveromyces marxianus |
| YMX003453 | 1.843 | Kluyveromyces marxianus | Kluyveromyces marxianus |
| YMX003264 | 1.841 | Kluyveromyces marxianus | Kluyveromyces marxianus |
| YMX003772 | 1.841 | Kluyveromyces marxianus | Kluyveromyces marxianus |
| YMX001077 | 1.84 | Kluyveromyces marxianus | Kluyveromyces marxianus |
| YMX001264 | 1.84 | Kluyveromyces marxianus | Kluyveromyces marxianus |
| YMX003250 | 1.84 | Kluyveromyces marxianus | Kluyveromyces marxianus |
| YMX003354 | 1.831 | Kluyveromyces marxianus | Kluyveromyces marxianus |
| YMX002331 | 1.83 | Kluyveromyces marxianus | Kluyveromyces marxianus |
| YMX003773 | 1.83 | Kluyveromyces marxianus | Kluyveromyces marxianus |
| YMX002005 | 1.823 | Kluyveromyces marxianus | Kluyveromyces marxianus |

| | | | |
|---|---|---|---|
| YMX004051 | 1.823 | Kluyveromyces marxianus | Kluyveromyces marxianus |
| YMX003027 | 1.813 | Kluyveromyces marxianus | Kluyveromyces marxianus |
| YMX003263 | 1.81 | Kluyveromyces marxianus | Kluyveromyces marxianus |
| YMX001999 | 1.8 | Kluyveromyces marxianus | Kluyveromyces marxianus |
| YMX003252 | 1.8 | Kluyveromyces marxianus | Kluyveromyces marxianus |
| YMX001256 | 1.792 | Kluyveromyces marxianus | Kluyveromyces marxianus |
| YMX002207 | 1.79 | Kluyveromyces marxianus | Kluyveromyces marxianus |
| YMX002333 | 1.79 | Kluyveromyces marxianus | Kluyveromyces marxianus |
| YMX003262 | 1.79 | Kluyveromyces marxianus | Kluyveromyces marxianus |
| YMX003350 | 1.79 | Kluyveromyces marxianus | Kluyveromyces marxianus |
| YMX003351 | 1.79 | Kluyveromyces marxianus | Kluyveromyces marxianus |
| YMX003672 | 1.79 | Kluyveromyces marxianus | Kluyveromyces marxianus |
| YMX002335 | 1.78 | Kluyveromyces marxianus | Kluyveromyces marxianus |
| YMX003345 | 1.77 | Kluyveromyces marxianus | Kluyveromyces marxianus |
| YMX002931 | 1.763 | Kluyveromyces marxianus | Kluyveromyces marxianus |
| YMX001654 | 1.76 | Kluyveromyces marxianus | Kluyveromyces marxianus |
| YMX002351 | 1.75 | Kluyveromyces marxianus | Kluyveromyces marxianus |
| YMX003344 | 1.742 | Kluyveromyces marxianus | Kluyveromyces marxianus |
| YMX003784 | 1.741 | Kluyveromyces marxianus | Kluyveromyces marxianus |
| YMX003342 | 1.74 | Kluyveromyces marxianus | Kluyveromyces marxianus |
| YMX003783 | 1.74 | Kluyveromyces marxianus | Kluyveromyces marxianus |
| YMX003785 | 1.74 | Kluyveromyces marxianus | Kluyveromyces marxianus |
| YMX003260 | 1.73 | Kluyveromyces marxianus | Kluyveromyces marxianus |
| YMX001122 | 1.713 | Kluyveromyces marxianus | Kluyveromyces marxianus |
| YMX001988 | 1.703 | Kluyveromyces marxianus | Kluyveromyces marxianus |
| YMX002006 | 1.68 | Kluyveromyces marxianus | Kluyveromyces marxianus |
| YMX002145 | 1.68 | Kluyveromyces marxianus | Kluyveromyces marxianus |
| YMX003340 | 1.68 | Kluyveromyces marxianus | Kluyveromyces marxianus |
| YMX003360 | 1.67 | Kluyveromyces marxianus | Kluyveromyces marxianus |
| YMX002495 | 1.61 | Kluyveromyces marxianus | Kluyveromyces marxianus |
| YMX001986 | 1.603 | Kluyveromyces marxianus | Kluyveromyces marxianus |
| YMX002340 | 1.591 | Kluyveromyces marxianus | Kluyveromyces marxianus |
| YMX002344 | 1.574 | Kluyveromyces marxianus | Kluyveromyces marxianus |
| YMX001881 | 1.57 | Kluyveromyces marxianus | Kluyveromyces marxianus |
| YMX002940 | 1.55 | Kluyveromyces marxianus | Kluyveromyces marxianus |
| YMX003452 | 1.54 | Kluyveromyces marxianus | Kluyveromyces marxianus |
| YMX002159 | 1.52 | Kluyveromyces marxianus | Kluyveromyces marxianus |
| YMX002149 | 1.491 | Kluyveromyces marxianus | Kluyveromyces marxianus |
| YMX003188 | 2.231 | Kluyveromyces marxianus | Pichia kluyveri |
| YMX002546 | 2.01 | Kluyveromyces marxianus | Pichia kluyveri |
| YMX002551 | 2.22 | Kluyveromyces marxianus | Pichia kudriavzevii |
| YMX002562 | 2.21 | Kluyveromyces marxianus | Pichia kudriavzevii |
| YMX002560 | 2.16 | Kluyveromyces marxianus | Pichia kudriavzevii |
| YMX002552 | 2.103 | Kluyveromyces marxianus | Pichia kudriavzevii |
| YMX002568 | 2.09 | Kluyveromyces marxianus | Pichia kudriavzevii |
| YMX002549 | 2.08 | Kluyveromyces marxianus | Pichia kudriavzevii |
| YMX002561 | 2.06 | Kluyveromyces marxianus | Pichia kudriavzevii |

| | | | |
|---|---|---|---|
| YMX002553 | 2.032 | Kluyveromyces marxianus | Pichia kudriavzevii |
| YMX003665 | 2.02 | Kluyveromyces marxianus | Pichia kudriavzevii |
| YMX003025 | 1.95 | Kluyveromyces marxianus | Pichia kudriavzevii |
| YMX003253 | 1.92 | Kluyveromyces marxianus | Pichia kudriavzevii |
| YMX002930 | 1.88 | Kluyveromyces marxianus | Pichia kudriavzevii |
| YMX001697 | 1.86 | Kluyveromyces marxianus | Pichia kudriavzevii |
| YMX002566 | 1.85 | Kluyveromyces marxianus | Pichia kudriavzevii |
| YMX002545 | 1.834 | Kluyveromyces marxianus | Pichia kudriavzevii |
| YMX003339 | 1.78 | Kluyveromyces marxianus | Pichia kudriavzevii |
| YMX003654 | 1.74 | Kluyveromyces marxianus | Pichia kudriavzevii |
| YMX003244 | 1.94 | Kluyveromyces marxianus | Pichia manshurica |
| YMX001649 | 1.72 | Kluyveromyces marxianus | Pichia manshurica |
| YMX003039 | 1.68 | Kluyveromyces marxianus | Pichia manshurica |
| YMX001587 | 1.8 | Kluyveromyces marxianus | Pichia membranifaciens |
| YMX001499 | 1.84 | Kluyveromyces marxianus | Saccharomyces cerevisiae |
| YMX003673 | 1.54 | Pichia kudriavzevii | Candida xylopsoci |
| YMX001903 | 2.11 | Pichia kudriavzevii | Fungal sp |
| YMX002617 | 2.161 | Pichia kudriavzevii | Kluyveromyces marxianus |
| YMX002631 | 2.16 | Pichia kudriavzevii | Kluyveromyces marxianus |
| YMX001594 | 2.282 | Pichia kudriavzevii | Pichia kudriavzevii |
| YMX002589 | 2.263 | Pichia kudriavzevii | Pichia kudriavzevii |
| YMX002993 | 2.26 | Pichia kudriavzevii | Pichia kudriavzevii |
| YMX003358 | 2.25 | Pichia kudriavzevii | Pichia kudriavzevii |
| YMX001709 | 2.24 | Pichia kudriavzevii | Pichia kudriavzevii |
| YMX000787 | 2.232 | Pichia kudriavzevii | Pichia kudriavzevii |
| YMX003176 | 2.231 | Pichia kudriavzevii | Pichia kudriavzevii |
| YMX003469 | 2.2 | Pichia kudriavzevii | Pichia kudriavzevii |
| YMX003574 | 2.18 | Pichia kudriavzevii | Pichia kudriavzevii |
| YMX001326 | 2.16 | Pichia kudriavzevii | Pichia kudriavzevii |
| YMX003593 | 2.16 | Pichia kudriavzevii | Pichia kudriavzevii |
| YMX001379 | 2.15 | Pichia kudriavzevii | Pichia kudriavzevii |
| YMX003504 | 2.15 | Pichia kudriavzevii | Pichia kudriavzevii |
| YMX002593 | 2.13 | Pichia kudriavzevii | Pichia kudriavzevii |
| YMX003343 | 2.13 | Pichia kudriavzevii | Pichia kudriavzevii |
| YMX002451 | 2.12 | Pichia kudriavzevii | Pichia kudriavzevii |
| YMX003744 | 2.12 | Pichia kudriavzevii | Pichia kudriavzevii |
| YMX001334 | 2.11 | Pichia kudriavzevii | Pichia kudriavzevii |
| YMX001329 | 2.1 | Pichia kudriavzevii | Pichia kudriavzevii |
| YMX001608 | 2.1 | Pichia kudriavzevii | Pichia kudriavzevii |
| YMX003516 | 2.1 | Pichia kudriavzevii | Pichia kudriavzevii |
| YMX002598 | 2.094 | Pichia kudriavzevii | Pichia kudriavzevii |
| YMX002979 | 2.09 | Pichia kudriavzevii | Pichia kudriavzevii |
| YMX001341 | 2.084 | Pichia kudriavzevii | Pichia kudriavzevii |
| YMX003764 | 2.078 | Pichia kudriavzevii | Pichia kudriavzevii |
| YMX002609 | 2.071 | Pichia kudriavzevii | Pichia kudriavzevii |
| YMX001899 | 2.07 | Pichia kudriavzevii | Pichia kudriavzevii |
| YMX001981 | 2.06 | Pichia kudriavzevii | Pichia kudriavzevii |

| | | | |
|---|---|---|---|
| YMX001253 | 2.05 | Pichia kudriavzevii | Pichia kudriavzevii |
| YMX002015 | 2.05 | Pichia kudriavzevii | Pichia kudriavzevii |
| YMX003514 | 2.05 | Pichia kudriavzevii | Pichia kudriavzevii |
| YMX002009 | 2.03 | Pichia kudriavzevii | Pichia kudriavzevii |
| YMX002980 | 2.03 | Pichia kudriavzevii | Pichia kudriavzevii |
| YMX003180 | 2.03 | Pichia kudriavzevii | Pichia kudriavzevii |
| YMX003449 | 2.022 | Pichia kudriavzevii | Pichia kudriavzevii |
| YMX001585 | 2.021 | Pichia kudriavzevii | Pichia kudriavzevii |
| YMX001323 | 2.02 | Pichia kudriavzevii | Pichia kudriavzevii |
| YMX001700 | 2.01 | Pichia kudriavzevii | Pichia kudriavzevii |
| YMX002453 | 2.01 | Pichia kudriavzevii | Pichia kudriavzevii |
| YMX001327 | 2.003 | Pichia kudriavzevii | Pichia kudriavzevii |
| YMX001318 | 2 | Pichia kudriavzevii | Pichia kudriavzevii |
| YMX002854 | 2 | Pichia kudriavzevii | Pichia kudriavzevii |
| YMX001718 | 1.99 | Pichia kudriavzevii | Pichia kudriavzevii |
| YMX001914 | 1.99 | Pichia kudriavzevii | Pichia kudriavzevii |
| YMX003924 | 1.98 | Pichia kudriavzevii | Pichia kudriavzevii |
| YMX001380 | 1.96 | Pichia kudriavzevii | Pichia kudriavzevii |
| YMX001372 | 1.954 | Pichia kudriavzevii | Pichia kudriavzevii |
| YMX003063 | 1.95 | Pichia kudriavzevii | Pichia kudriavzevii |
| YMX003168 | 1.95 | Pichia kudriavzevii | Pichia kudriavzevii |
| YMX003346 | 1.95 | Pichia kudriavzevii | Pichia kudriavzevii |
| YMX001390 | 1.942 | Pichia kudriavzevii | Pichia kudriavzevii |
| YMX001371 | 1.94 | Pichia kudriavzevii | Pichia kudriavzevii |
| YMX002584 | 1.93 | Pichia kudriavzevii | Pichia kudriavzevii |
| YMX001241 | 1.92 | Pichia kudriavzevii | Pichia kudriavzevii |
| YMX001373 | 1.92 | Pichia kudriavzevii | Pichia kudriavzevii |
| YMX003149 | 1.92 | Pichia kudriavzevii | Pichia kudriavzevii |
| YMX003336 | 1.91 | Pichia kudriavzevii | Pichia kudriavzevii |
| YMX003684 | 1.89 | Pichia kudriavzevii | Pichia kudriavzevii |
| YMX002436 | 1.88 | Pichia kudriavzevii | Pichia kudriavzevii |
| YMX001992 | 1.87 | Pichia kudriavzevii | Pichia kudriavzevii |
| YMX002432 | 1.86 | Pichia kudriavzevii | Pichia kudriavzevii |
| YMX003677 | 1.854 | Pichia kudriavzevii | Pichia kudriavzevii |
| YMX001977 | 1.844 | Pichia kudriavzevii | Pichia kudriavzevii |
| YMX003668 | 1.821 | Pichia kudriavzevii | Pichia kudriavzevii |
| YMX003019 | 1.82 | Pichia kudriavzevii | Pichia kudriavzevii |
| YMX003312 | 1.793 | Pichia kudriavzevii | Pichia kudriavzevii |
| YMX001321 | 1.76 | Pichia kudriavzevii | Pichia kudriavzevii |
| YMX002840 | 1.723 | Pichia kudriavzevii | Pichia kudriavzevii |
| YMX003623 | 1.98 | Pichia kudriavzevii | Pichia manshurica |
| YMX002635 | 2.32 | Pichia kudriavzevii | Saccharomyces cerevisiae |
| YMX002599 | 2.31 | Pichia kudriavzevii | Saccharomyces cerevisiae |
| YMX002622 | 2.26 | Pichia kudriavzevii | Saccharomyces cerevisiae |
| YMX001734 | 2.12 | Pichia kudriavzevii | Saccharomyces cerevisiae |
| YMX002419 | 2.11 | Pichia kudriavzevii | Saccharomyces cerevisiae |
| YMX002417 | 2.023 | Pichia kudriavzevii | Saccharomyces cerevisiae |

| | | | |
|---|---|---|---|
| YMX003070 | 1.97 | Pichia kudriavzevii | Saccharomyces cerevisiae |
| YMX003953 | 1.93 | Pichia kudriavzevii | Saccharomyces cerevisiae |
| YMX001349 | 1.83 | Pichia kudriavzevii | Saccharomyces cerevisiae |
| YMX001737 | 1.79 | Pichia kudriavzevii | Saccharomyces cerevisiae |
| YMX000702 | 2.08 | Pichia manshurica | Kluyveromyces marxianus |
| YMX001177 | 1.85 | Pichia manshurica | Kluyveromyces marxianus |
| YMX000641 | 2.09 | Pichia manshurica | Pichia kluyveri |
| YMX000959 | 1.724 | Pichia manshurica | Pichia kluyveri |
| YMX002407 | 2.26 | Pichia manshurica | Pichia kudriavzevii |
| YMX002588 | 2.21 | Pichia manshurica | Pichia kudriavzevii |
| YMX003567 | 2.17 | Pichia manshurica | Pichia kudriavzevii |
| YMX001303 | 2.043 | Pichia manshurica | Pichia kudriavzevii |
| YMX002571 | 1.991 | Pichia manshurica | Pichia kudriavzevii |
| YMX001306 | 1.97 | Pichia manshurica | Pichia kudriavzevii |
| YMX001703 | 1.9 | Pichia manshurica | Pichia kudriavzevii |
| YMX003664 | 1.87 | Pichia manshurica | Pichia kudriavzevii |
| YMX003564 | 1.74 | Pichia manshurica | Pichia kudriavzevii |
| YMX004013 | 2.26 | Pichia manshurica | Pichia manshurica |
| YMX004012 | 2.22 | Pichia manshurica | Pichia manshurica |
| YMX002943 | 2.2 | Pichia manshurica | Pichia manshurica |
| YMX002944 | 2.2 | Pichia manshurica | Pichia manshurica |
| YMX002755 | 2.17 | Pichia manshurica | Pichia manshurica |
| YMX002757 | 2.17 | Pichia manshurica | Pichia manshurica |
| YMX002410 | 2.16 | Pichia manshurica | Pichia manshurica |
| YMX002737 | 2.16 | Pichia manshurica | Pichia manshurica |
| YMX003751 | 2.154 | Pichia manshurica | Pichia manshurica |
| YMX000637 | 2.14 | Pichia manshurica | Pichia manshurica |
| YMX002935 | 2.12 | Pichia manshurica | Pichia manshurica |
| YMX002641 | 2.114 | Pichia manshurica | Pichia manshurica |
| YMX000648 | 2.07 | Pichia manshurica | Pichia manshurica |
| YMX000725 | 2.06 | Pichia manshurica | Pichia manshurica |
| YMX002933 | 2.06 | Pichia manshurica | Pichia manshurica |
| YMX004059 | 2 | Pichia manshurica | Pichia manshurica |
| YMX001950 | 1.99 | Pichia manshurica | Pichia manshurica |
| YMX002473 | 1.99 | Pichia manshurica | Pichia manshurica |
| YMX002001 | 1.98 | Pichia manshurica | Pichia manshurica |
| YMX003660 | 1.98 | Pichia manshurica | Pichia manshurica |
| YMX004009 | 1.96 | Pichia manshurica | Pichia manshurica |
| YMX002923 | 1.95 | Pichia manshurica | Pichia manshurica |
| YMX000643 | 1.94 | Pichia manshurica | Pichia manshurica |
| YMX000697 | 1.924 | Pichia manshurica | Pichia manshurica |
| YMX003656 | 1.92 | Pichia manshurica | Pichia manshurica |
| YMX003535 | 1.91 | Pichia manshurica | Pichia manshurica |
| YMX003657 | 1.91 | Pichia manshurica | Pichia manshurica |
| YMX003347 | 1.882 | Pichia manshurica | Pichia manshurica |
| YMX002994 | 1.88 | Pichia manshurica | Pichia manshurica |
| YMX002977 | 1.87 | Pichia manshurica | Pichia manshurica |

| | | | |
|---|---|---|---|
| YMX001651 | 1.86 | Pichia manshurica | Pichia manshurica |
| YMX003536 | 1.86 | Pichia manshurica | Pichia manshurica |
| YMX003671 | 1.86 | Pichia manshurica | Pichia manshurica |
| YMX002957 | 1.85 | Pichia manshurica | Pichia manshurica |
| YMX002364 | 1.82 | Pichia manshurica | Pichia manshurica |
| YMX003760 | 1.811 | Pichia manshurica | Pichia manshurica |
| YMX002363 | 1.81 | Pichia manshurica | Pichia manshurica |
| YMX002508 | 1.8 | Pichia manshurica | Pichia manshurica |
| YMX001962 | 1.78 | Pichia manshurica | Pichia manshurica |
| YMX003341 | 1.734 | Pichia manshurica | Pichia manshurica |
| YMX003338 | 1.73 | Pichia manshurica | Pichia manshurica |
| YMX003767 | 1.714 | Pichia manshurica | Pichia manshurica |
| YMX001028 | 2.01 | Pichia manshurica | Pichia membranifaciens |
| YMX001030 | 1.73 | Pichia manshurica | Pichia membranifaciens |
| YMX002749 | 2.16 | Pichia manshurica | Pichia sp |
| YMX001533 | 2.15 | Pichia manshurica | Pichia sp |
| YMX001693 | 2.1 | Pichia manshurica | Pichia sp |
| YMX003549 | 2.06 | Pichia manshurica | Pichia sp |
| YMX001504 | 2.02 | Pichia manshurica | Pichia sp |
| YMX002324 | 2.001 | Pichia manshurica | Pichia sp |
| YMX001785 | 1.982 | Pichia manshurica | Pichia sp |
| YMX001525 | 1.944 | Pichia manshurica | Pichia sp |
| YMX001780 | 1.924 | Pichia manshurica | Pichia sp |
| YMX001536 | 1.92 | Pichia manshurica | Pichia sp |
| YMX001787 | 1.823 | Pichia manshurica | Pichia sp |
| YMX000949 | 1.74 | Pichia manshurica | Pichia sp |
| YMX003061 | 1.92 | Pichia manshurica | Saccharomyces cerevisiae |
| YMX003069 | 1.91 | Pichia manshurica | Saccharomyces cerevisiae |
| YMX001657 | 1.9 | Pichia manshurica | Saccharomyces cerevisiae |
| YMX000163 | 1.863 | Saccharomyces cerevisiae | Candida ethanolica |
| YMX000242 | 2.01 | Saccharomyces cerevisiae | Kluyveromyces marxianus |
| YMX000130 | 1.821 | Saccharomyces cerevisiae | Penicillium simplicissimum |
| YMX003801 | 1.76 | Saccharomyces cerevisiae | Pichia kluyveri |
| YMX001153 | 2.17 | Saccharomyces cerevisiae | Saccharomyces cerevisiae |
| YMX000518 | 2.127 | Saccharomyces cerevisiae | Saccharomyces cerevisiae |
| YMX002905 | 2.121 | Saccharomyces cerevisiae | Saccharomyces cerevisiae |
| YMX000546 | 2.114 | Saccharomyces cerevisiae | Saccharomyces cerevisiae |
| YMX003506 | 2.111 | Saccharomyces cerevisiae | Saccharomyces cerevisiae |
| YMX000937 | 2.104 | Saccharomyces cerevisiae | Saccharomyces cerevisiae |
| YMX002809 | 2.081 | Saccharomyces cerevisiae | Saccharomyces cerevisiae |
| YMX002461 | 2.06 | Saccharomyces cerevisiae | Saccharomyces cerevisiae |
| YMX001922 | 2.05 | Saccharomyces cerevisiae | Saccharomyces cerevisiae |
| YMX000330 | 2.04 | Saccharomyces cerevisiae | Saccharomyces cerevisiae |
| YMX000673 | 2.03 | Saccharomyces cerevisiae | Saccharomyces cerevisiae |
| YMX000818 | 2.013 | Saccharomyces cerevisiae | Saccharomyces cerevisiae |
| YMX001684 | 2.002 | Saccharomyces cerevisiae | Saccharomyces cerevisiae |
| YMX003194 | 2.002 | Saccharomyces cerevisiae | Saccharomyces cerevisiae |

| | | | |
|---|---|---|---|
| YMX000108 | 2.001 | Saccharomyces cerevisiae | Saccharomyces cerevisiae |
| YMX003402 | 2.001 | Saccharomyces cerevisiae | Saccharomyces cerevisiae |
| YMX002573 | 1.99 | Saccharomyces cerevisiae | Saccharomyces cerevisiae |
| YMX001951 | 1.98 | Saccharomyces cerevisiae | Saccharomyces cerevisiae |
| YMX000369 | 1.975 | Saccharomyces cerevisiae | Saccharomyces cerevisiae |
| YMX002585 | 1.974 | Saccharomyces cerevisiae | Saccharomyces cerevisiae |
| YMX000963 | 1.973 | Saccharomyces cerevisiae | Saccharomyces cerevisiae |
| YMX000306 | 1.972 | Saccharomyces cerevisiae | Saccharomyces cerevisiae |
| YMX001393 | 1.971 | Saccharomyces cerevisiae | Saccharomyces cerevisiae |
| YMX000594 | 1.97 | Saccharomyces cerevisiae | Saccharomyces cerevisiae |
| YMX000841 | 1.97 | Saccharomyces cerevisiae | Saccharomyces cerevisiae |
| YMX000988 | 1.97 | Saccharomyces cerevisiae | Saccharomyces cerevisiae |
| YMX002401 | 1.97 | Saccharomyces cerevisiae | Saccharomyces cerevisiae |
| YMX002479 | 1.97 | Saccharomyces cerevisiae | Saccharomyces cerevisiae |
| YMX004014 | 1.97 | Saccharomyces cerevisiae | Saccharomyces cerevisiae |
| YMX000743 | 1.964 | Saccharomyces cerevisiae | Saccharomyces cerevisiae |
| YMX000721 | 1.96 | Saccharomyces cerevisiae | Saccharomyces cerevisiae |
| YMX000935 | 1.96 | Saccharomyces cerevisiae | Saccharomyces cerevisiae |
| YMX001298 | 1.96 | Saccharomyces cerevisiae | Saccharomyces cerevisiae |
| YMX003313 | 1.96 | Saccharomyces cerevisiae | Saccharomyces cerevisiae |
| YMX000361 | 1.959 | Saccharomyces cerevisiae | Saccharomyces cerevisiae |
| YMX000284 | 1.953 | Saccharomyces cerevisiae | Saccharomyces cerevisiae |
| YMX002449 | 1.953 | Saccharomyces cerevisiae | Saccharomyces cerevisiae |
| YMX000194 | 1.952 | Saccharomyces cerevisiae | Saccharomyces cerevisiae |
| YMX002450 | 1.952 | Saccharomyces cerevisiae | Saccharomyces cerevisiae |
| YMX003625 | 1.95 | Saccharomyces cerevisiae | Saccharomyces cerevisiae |
| YMX000151 | 1.94 | Saccharomyces cerevisiae | Saccharomyces cerevisiae |
| YMX001733 | 1.94 | Saccharomyces cerevisiae | Saccharomyces cerevisiae |
| YMX002521 | 1.94 | Saccharomyces cerevisiae | Saccharomyces cerevisiae |
| YMX003002 | 1.94 | Saccharomyces cerevisiae | Saccharomyces cerevisiae |
| YMX002185 | 1.932 | Saccharomyces cerevisiae | Saccharomyces cerevisiae |
| YMX000362 | 1.93 | Saccharomyces cerevisiae | Saccharomyces cerevisiae |
| YMX002458 | 1.93 | Saccharomyces cerevisiae | Saccharomyces cerevisiae |
| YMX002665 | 1.93 | Saccharomyces cerevisiae | Saccharomyces cerevisiae |
| YMX003553 | 1.93 | Saccharomyces cerevisiae | Saccharomyces cerevisiae |
| YMX000232 | 1.92 | Saccharomyces cerevisiae | Saccharomyces cerevisiae |
| YMX001036 | 1.92 | Saccharomyces cerevisiae | Saccharomyces cerevisiae |
| YMX000116 | 1.914 | Saccharomyces cerevisiae | Saccharomyces cerevisiae |
| YMX000865 | 1.912 | Saccharomyces cerevisiae | Saccharomyces cerevisiae |
| YMX001949 | 1.911 | Saccharomyces cerevisiae | Saccharomyces cerevisiae |
| YMX001566 | 1.91 | Saccharomyces cerevisiae | Saccharomyces cerevisiae |
| YMX002642 | 1.9 | Saccharomyces cerevisiae | Saccharomyces cerevisiae |
| YMX003962 | 1.9 | Saccharomyces cerevisiae | Saccharomyces cerevisiae |
| YMX000018 | 1.892 | Saccharomyces cerevisiae | Saccharomyces cerevisiae |
| YMX000219 | 1.89 | Saccharomyces cerevisiae | Saccharomyces cerevisiae |
| YMX002071 | 1.872 | Saccharomyces cerevisiae | Saccharomyces cerevisiae |
| YMX001756 | 1.87 | Saccharomyces cerevisiae | Saccharomyces cerevisiae |

| | | | |
|---|---|---|---|
| YMX000577 | 1.869 | Saccharomyces cerevisiae | Saccharomyces cerevisiae |
| YMX001132 | 1.863 | Saccharomyces cerevisiae | Saccharomyces cerevisiae |
| YMX001225 | 1.86 | Saccharomyces cerevisiae | Saccharomyces cerevisiae |
| YMX002190 | 1.86 | Saccharomyces cerevisiae | Saccharomyces cerevisiae |
| YMX000626 | 1.84 | Saccharomyces cerevisiae | Saccharomyces cerevisiae |
| YMX002497 | 1.83 | Saccharomyces cerevisiae | Saccharomyces cerevisiae |
| YMX003051 | 1.83 | Saccharomyces cerevisiae | Saccharomyces cerevisiae |
| YMX002020 | 1.824 | Saccharomyces cerevisiae | Saccharomyces cerevisiae |
| YMX000188 | 1.82 | Saccharomyces cerevisiae | Saccharomyces cerevisiae |
| YMX001658 | 1.82 | Saccharomyces cerevisiae | Saccharomyces cerevisiae |
| YMX002378 | 1.82 | Saccharomyces cerevisiae | Saccharomyces cerevisiae |
| YMX003889 | 1.82 | Saccharomyces cerevisiae | Saccharomyces cerevisiae |
| YMX003265 | 1.811 | Saccharomyces cerevisiae | Saccharomyces cerevisiae |
| YMX001273 | 1.803 | Saccharomyces cerevisiae | Saccharomyces cerevisiae |
| YMX002041 | 1.801 | Saccharomyces cerevisiae | Saccharomyces cerevisiae |
| YMX001191 | 1.8 | Saccharomyces cerevisiae | Saccharomyces cerevisiae |
| YMX002713 | 1.8 | Saccharomyces cerevisiae | Saccharomyces cerevisiae |
| YMX001849 | 1.79 | Saccharomyces cerevisiae | Saccharomyces cerevisiae |
| YMX002468 | 1.79 | Saccharomyces cerevisiae | Saccharomyces cerevisiae |
| YMX001874 | 1.78 | Saccharomyces cerevisiae | Saccharomyces cerevisiae |
| YMX002427 | 1.78 | Saccharomyces cerevisiae | Saccharomyces cerevisiae |
| YMX003390 | 1.771 | Saccharomyces cerevisiae | Saccharomyces cerevisiae |
| YMX003433 | 1.763 | Saccharomyces cerevisiae | Saccharomyces cerevisiae |
| YMX001084 | 1.762 | Saccharomyces cerevisiae | Saccharomyces cerevisiae |
| YMX001345 | 1.76 | Saccharomyces cerevisiae | Saccharomyces cerevisiae |
| YMX001901 | 1.76 | Saccharomyces cerevisiae | Saccharomyces cerevisiae |
| YMX002332 | 1.76 | Saccharomyces cerevisiae | Saccharomyces cerevisiae |
| YMX004105 | 1.76 | Saccharomyces cerevisiae | Saccharomyces cerevisiae |
| YMX001011 | 1.753 | Saccharomyces cerevisiae | Saccharomyces cerevisiae |
| YMX003073 | 1.752 | Saccharomyces cerevisiae | Saccharomyces cerevisiae |
| YMX001514 | 1.75 | Saccharomyces cerevisiae | Saccharomyces cerevisiae |
| YMX002452 | 1.74 | Saccharomyces cerevisiae | Saccharomyces cerevisiae |
| YMX003398 | 1.74 | Saccharomyces cerevisiae | Saccharomyces cerevisiae |
| YMX003170 | 1.73 | Saccharomyces cerevisiae | Saccharomyces cerevisiae |
| YMX003461 | 1.73 | Saccharomyces cerevisiae | Saccharomyces cerevisiae |
| YMX002833 | 1.72 | Saccharomyces cerevisiae | Saccharomyces cerevisiae |
| YMX002547 | 1.7 | Saccharomyces cerevisiae | Saccharomyces cerevisiae |
| YMX003409 | 1.7 | Saccharomyces cerevisiae | Saccharomyces cerevisiae |
| YMX000012 | 1.688 | Saccharomyces cerevisiae | Saccharomyces cerevisiae |
| YMX003697 | 1.68 | Saccharomyces cerevisiae | Saccharomyces cerevisiae |
| YMX001969 | 1.584 | Saccharomyces cerevisiae | Saccharomyces cerevisiae |
| YMX001945 | 1.58 | Saccharomyces cerevisiae | Saccharomyces cerevisiae |
| YMX002466 | 1.504 | Saccharomyces cerevisiae | Saccharomyces cerevisiae |
| YMX001586 | 1.93 | Saccharomyces cerevisiae | Saccharomyces kudriavzevii |
| YMX000557 | 2.07 | Saccharomyces cerevisiae | Saccharomyces paradoxus |
| YMX002690 | 2.02 | Saccharomyces cerevisiae | Saccharomyces paradoxus |
| YMX000649 | 1.914 | Saccharomyces cerevisiae | Saccharomyces paradoxus |

| | | | |
|---|---|---|---|
| YMX002953 | 1.91 | Saccharomyces cerevisiae | Saccharomyces paradoxus |
| YMX000465 | 1.89 | Saccharomyces cerevisiae | Saccharomyces paradoxus |
| YMX003824 | 1.86 | Saccharomyces cerevisiae | Saccharomyces paradoxus |
| YMX001537 | 1.78 | Saccharomyces cerevisiae | Saccharomyces paradoxus |
| YMX000578 | 1.777 | Saccharomyces cerevisiae | Saccharomyces paradoxus |
| YMX002305 | 1.54 | Saccharomyces cerevisiae | Saccharomyces paradoxus |

**Table S2.** Assembly statistics of four *Kazachstania humilis* strains from agave fermentation and the CLIB 1323 type strain

| | Agave fermentation strains (YMX.1.0) | | | | Type strain |
|---|---|---|---|---|---|
| | **YMX000387*** | **YMX000554*** | **YMX003162*** | **YMX004033** | **CLIB 1323** |
| No. scaffolds | 194 | 239 | 256 | 21 | 16 |
| Largest scaffold | 2,388,392 | 2,389,372 | 2,383,625 | 2,288,783 | 2,314,217 |
| Total length | 17,384,806 | 19,010,241 | 19,113,864 | 13,956,217 | 13,969,787 |
| GC (%) | 49.16 | 49.23 | 49.3 | 49.16 | 49.15 |
| N50 | 975,232 | 957,909 | 863,393 | 1,026,251 | 1,009,204 |
| N75 | 377,545 | 270,810 | 176,989 | 737,073 | 934,176 |
| L50 | 7 | 8 | 8 | 6 | 5 |
| L75 | 14 | 16 | 17 | 10 | 8 |
| No. genes | 5,404 | 5,452 | 5,463 | 5,016 | 4,295 |
| No. Ns | 219,600 | 192,500 | 199,900 | 0 | 0 |
| No. Ns /100 kb | 1,263.17 | 1,012.61 | 1,045.84 | 0 | 0 |

* This study

**Table S3.** Summary of SyRI structural annotation of Kazachstania humilis genomes (PDF file)

| **Variant type** | **Count** | **CLIB1323 Length (bp)** | **YMX004033 Length (bp)** |
|---|---|---|---|
| Syntenic regions | 197 | 11,840,890 | 11,888,173 |
| Inversions | 2 | 3,673 | 3,958 |
| Translocations | 66 | 2,305,434 | 2,298,330 |
| Duplications in CLIB1323 | 137 | 373,151 | - |
| Duplications in YMX004033 | 71 | - | 205,241 |
| Not aligned in CLIB1323 | 182 | 244,442 | - |
| Not aligned in YMX004033 | 169 | - | 692,968 |
| SNPs | 503,408 | 503,408 | 503,408 |
| Insertions | 33,111 | - | 90,355 |
| Deletions | 23,952 | 79,463 | - |
| Copy gains in YMX004033 | 26 | - | 44,120 |
| Copy losses in CLIB1323 | 24 | 66,450 | - |
| Highly diverged | 246 | 250,702 | 306,570 |
| Tandem repeats | 4 | 9,894 | 8,189 |
